# Antimicrobial activity of iron-depriving pyoverdines against human opportunistic pathogens

**DOI:** 10.1101/2023.07.18.549568

**Authors:** Vera Vollenweider, Karoline Rehm, Clara Chepkirui, Manuela Pérez-Berlanga, Magdalini Polymenidou, Jörn Piel, Laurent Bigler, Rolf Kümmerli

**Author notes:** **Corresponding author**: Rolf Kümmerli, Department of Quantitative Biomedicine, University of Zurich, Winterthurerstrasse 190, 8057 Zurich, Switzerland. /.

## Abstract

The global rise of antibiotic resistance calls for new drugs against bacterial pathogens. A common approach is to search for natural compounds deployed by microbes to inhibit competitors. Here we show that the iron chelating pyoverdines, siderophores produced by environmental *Pseudomonas* spp., have strong antibacterial properties by inducing iron starvation and growth arrest in pathogens. A screen of 320 natural *Pseudomonas* isolates used against 12 human pathogens uncovered several pyoverdines with particularly high antibacterial properties and distinct chemical characteristics. The most potent pyoverdine effectively reduced growth of the pathogens *Acinetobacter baumannii*, *Klebsiella pneumoniae* and *Staphylococcus aureus* in a concentration- and iron-dependent manner. Pyoverdine increased survival of infected *Galleria mellonella* host larvae, and showed low toxicity for the host, mammalian cell lines, and erythrocytes. Furthermore, experimental evolution combined with whole-genome sequencing revealed reduced potentials for resistance evolution compared to an antibiotic. Thus, pyoverdines from environmental strains have the potential to become a new class of sustainable antibacterials against specific human pathogens.

## Introduction

There are approximately 4.95 million fatalities associated with antibiotic-resistant bacteria worldwide each year^1^. This high mortality rate contrasts with the low development rate of new antibacterial agents for clinical applications^2,3^. Thus, the development of alternative therapies with higher sustainability to counteract the rapidly dwindling treatment options due to resistance evolution is imperative^4–6^. Numerous alternative approaches have been proposed including phage therapy, antimicrobial peptides and nanobodies, anti-virulence treatments to disarm pathogens, treatments that capitalize on ecological competition between susceptible and resistant bacteria, and the search for new antibacterial compounds produced by environmental microbes^7–12^.

Mining natural microbial communities has become a promising endeavour in the hunt for novel antimicrobials because most microbes deploy bioactive compounds to contend with their competitors in the diverse assemblies they live in. Competition often involves the secretion of secondary metabolites with specific antimicrobial properties against other microbes^13^. The isolation, characterization and synthesis of such secondary metabolites can result in novel prospective antibiotics for clinical applications^14,15^. Secondary metabolites that act as toxins to kill competitors are often considered promising candidates. However, microbes compete through a variety of mechanisms other than toxins. For example, there is increasing evidence that competition for iron is a main determinant of species interactions in soil and freshwater communities^16,17^. Competition for iron involves the secretion of siderophores, a class of secondary metabolites that scavenges environmental iron with high affinity^18^. Because siderophores are often species specific, they have two opposing effects: they make iron available for related community members possessing the matching receptor but withhold iron from competitors with non-matching receptors. Thus, siderophores can be competitive agents in inter-species interactions^19–22^.

Here, we apply the concept of siderophore-mediated iron competition observed in natural communities to human opportunistic pathogens. Specifically, we investigate whether siderophores from non-pathogenic environmental bacteria can induce iron starvation and growth arrest in human pathogens. We focus on pyoverdines, a class of siderophores with high iron affinity that are produced and secreted by fluorescent *Pseudomonas* spp. under iron-limited conditions^23^. Upon complexation with ferric iron, the iron-loaded pyoverdine is imported into the cell by receptors with high specificity and subsequently reduced to the bio-available ferrous iron^24,25^. Pyoverdines show an extraordinary structural diversity with more than 70 described variants that differ in their peptide backbone^26,27^, and have an extremely high affinity for iron (K_a_ = 10^32^ M^-1^, pyoverdine produced by *Pseudomonas fluorescens* biotype B)^28^. For these reasons, we propose that pyoverdines from non-pathogenic *Pseudomonas* spp. could be potent agents to reduce the growth of opportunistic human pathogens by intensifying iron competition.

To test our hypothesis, we screened pyoverdines from a library of 320 environmental *Pseudomonas* strains, isolated from soil and freshwater habitats, for their activity against 12 strains of human opportunistic bacteria. For the most promising pyoverdine candidates, we elucidated the chemical structure to assess diversity and to understand the chemical properties important for iron competition. Subsequently, we assessed the efficacy of the three top pyoverdine candidates against four human opportunistic pathogens (*Acinetobacter baumannii*, *Klebsiella pneumoniae*, *Pseudomonas aeruginosa* and *Staphylococcus aureus*) *in vitro* and/or *in vivo* via infections of the Greater wax moth larvae (*Galleria mellonella*). We further assessed their toxicity in the host, towards two mammalian cell lines and red blood cells. Finally, we used experimental evolution combined with whole-genome sequencing to study the potential of pathogens evolving resistance against pyoverdine treatment.

## Results

### Pyoverdines can inhibit the growth of human opportunistic pathogens

To assess the antibacterial properties of pyoverdines, we performed a growth-inhibition screen using a well-characterized collection of 320 natural *Pseudomonas* isolates^16,29–31^. The isolates stem from freshwater and soil communities and belong to the group of fluorescent pseudomonads (e.g, *P. fluorescens*, *P. putida*, *P. syringae*, *P. chlororaphis*) that are non-pathogenic to humans and can produce and secrete pyoverdines, a group of high iron-affinity siderophores^32^. Important to note is that several isolates are likely to produce secondary siderophores in addition to pyoverdine^33^. Given that secondary siderophores generally have lower iron affinity than pyoverdine, we anticipate treatment effects to be primarily driven by pyoverdines.

In a first screen, we evaluated how the sterile pyoverdine-containing supernatants of the 320 isolates affect the growth of 12 human opportunistic pathogens (Table 1). Specifically, we grew the pathogens in 70% fresh casamino acid medium (CAA) supplemented with 30% spent supernatant. We found high variation in the supernatant effect, ranging from complete pathogen growth inhibition to growth promotion compared to the control treatment (70% CAA supplemented with 30% sodium chloride 0.8% solution) (Figure 1A and Figure S1A). The mean supernatant effect across all pathogens showed a bimodal distribution with two peaks at 0.47 – 0.58 (approximately 50% growth inhibition) and 1.00 – 1.10 (no growth inhibition) (Figure S1B). Our observations are consistent with previous studies, in which supernatants containing siderophores can have both growth inhibitory and stimulatory effects^20,29^. For each supernatant, we quantified the number of pathogens that experienced a growth reduction of at least 20% compared to the control and found that 25 supernatants inhibited all 12 pathogens (Figure 1A and S1C).

**Figure 1.**
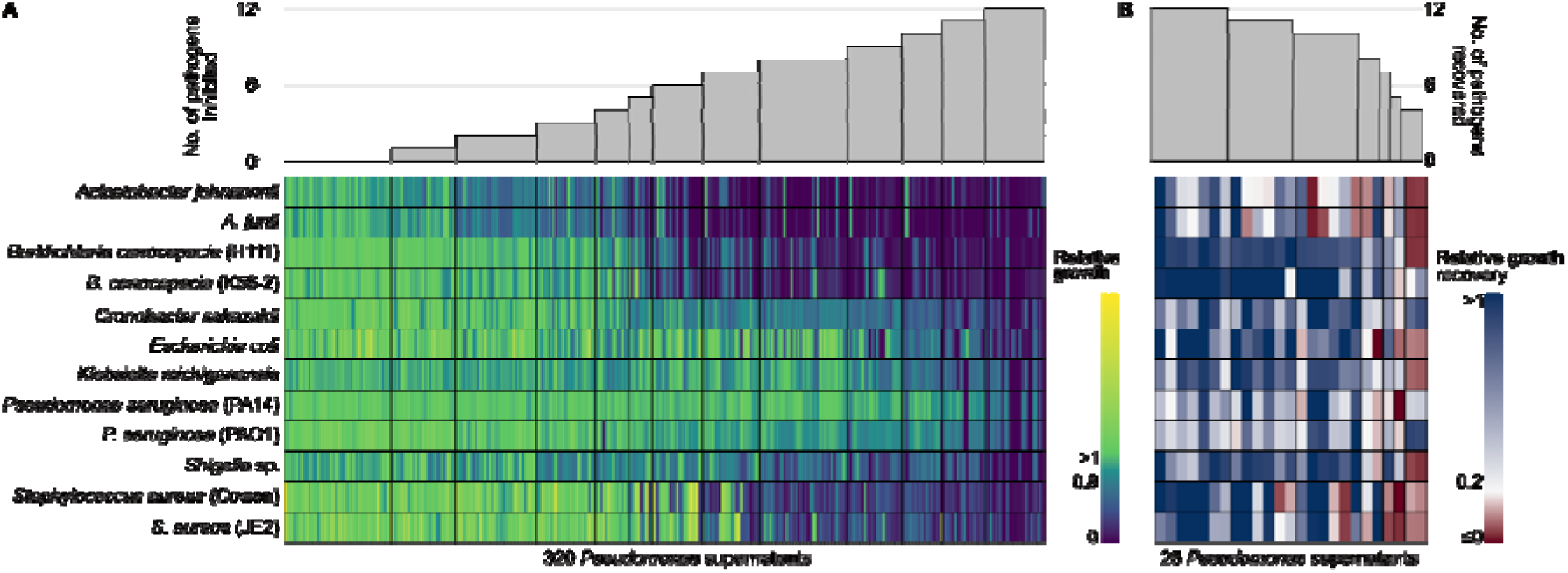
Effect of supernatants from environmental *Pseudomonas* isolates on the growth of 12 human opportunistic pathogens. (A) Screen to assess the extent to which pyoverdine-containing supernatants from 320 natural *Pseudomonas* isolates inhibit the growth of 12 human opportunistic pathogens. The heatmap depicts relative pathogen growth in the supernatant treatment (70% CAA + 30% spent supernatant) relative to the control (70% CAA + 30% sodium chloride solution), with values ranging from stimulation (yellow) to inhibition (blue) based on four independent replicates. Grey bars above the heatmap show the number of pathogens that were at least 20% inhibited in their growth by a given supernatant. The screen returned 25 supernatant candidates that inhibited the growth of all pathogens. (B) Control screen with the 25 top supernatant candidates to check whether pyoverdine causes the observed growth inhibition. The heatmap depicts the level of growth recovery in the 12 pathogens when iron was added to the supernatant, with values ranging from no recovery (red) to full recovery (blue). Growth recovery in iron-rich medium indicates that pyoverdines are involved in growth inhibition in iron-limited medium. Grey bars above the heatmap show the number of pathogens that experienced a relative growth recovery of at least 0.2. The screen returned 7 supernatant top candidates for which growth recovery occurred for all pathogens under iron-rich conditions.

**Table 1.** Species and strains of human opportunistic pathogens used for the pyoverdine growth inhibition assay.

| Species and strain name | Class | Gram-staining | Frequency of inhibiting supernatants |
| --- | --- | --- | --- |
| *Acinetobacter johnsonii* | Gammaproteobacteria | negative | 75.0 |
| *Acinetobacter junii* | Gammaproteobacteria | negative | 73.1 |
| *Burkholderia cenocepacia* H111 | Betaproteobacteria | negative | 52.2 |
| *Burkholderia cenocepacia* K56-2 | Betaproteobacteria | negative | 54.7 |
| *Cronobacter sakazakii* | Gammaproteobacteria | negative | 48.8 |
| *Escherichia coli* K12 | Gammaproteobacteria | negative | 31.6 |
| *Klebsiella michiganensis* | Gammaproteobacteria | negative | 30.9 |
| *Pseudomonas aeruginosa* PA14 | Gammaproteobacteria | negative | 9.7 |
| *Pseudomonas aeruginosa* PAO1 | Gammaproteobacteria | negative | 20.9 |
| *Shigella* sp. | Gammaproteobacteria | negative | 57.8 |
| *Staphylococcus aureus* Cowan | Bacilli | positive | 47.8 |
| *Staphylococcus aureus* JE2 | Bacilli | positive | 41.6 |

To confirm that the growth-inhibiting properties of the 25 most potent supernatants are indeed caused by pyoverdines and not by other agents in the supernatant, we repeated the above assay, but this time supplemented the supernatant with 40 µM FeCl_3_. Under such iron-replete conditions, pyoverdines (and secondary siderophores) should not be able to substantially sequester environmental iron, and pathogen growth should therefore be restored. We identified seven top candidate supernatants for which a relative growth recovery of at least 0.2 occurred for all 12 pathogens (Figure 1B). Among the remaining 18 supernatants, the overall rate of growth recovery was also high, indicating that pyoverdines are responsible for pathogen growth inhibition under iron-limited conditions in most cases.

Finally, we assessed the relationship between growth inhibitory effects of supernatants and the phylogenetic affiliation of the producing strains (Figure S2). We found that growth inhibitory effects were not restricted to a few closely related strains but were scattered across the phylogenetic tree. Nonetheless, there was a certain level of clustering with two *Pseudomonas* spp. clades showing an accumulation of strains with growth-inhibitory effects on pathogens. This analysis suggests that closely as well as distantly related strains can produce inhibitory pyoverdines.

### Growth-inhibiting pyoverdines show distinct chemical features

In previous studies, we developed a new method for pyoverdine structure elucidation from low-volume crude extracts using ultra-high-performance liquid chromatography high-resolution tandem mass spectrometry (UHPLC-HR-MS/MS)^27,34^. These studies included the pyoverdines extracted from the seven top candidate supernatants and the analysis therein yielded five structurally different pyoverdines (Figure 2A). We found one novel pyoverdine type (**1**, from isolate 3A06) and four pyoverdine types already known from the literature (**2**-**5**). The pyoverdine from isolate 3G07 can occur in either a linear or cyclic conformation (**2a** and **2b**). Three isolates (s3b09, s3b10, s3b12), all originating from the same soil sample, had an identical pyoverdine type (**3**).

**Figure 2.**
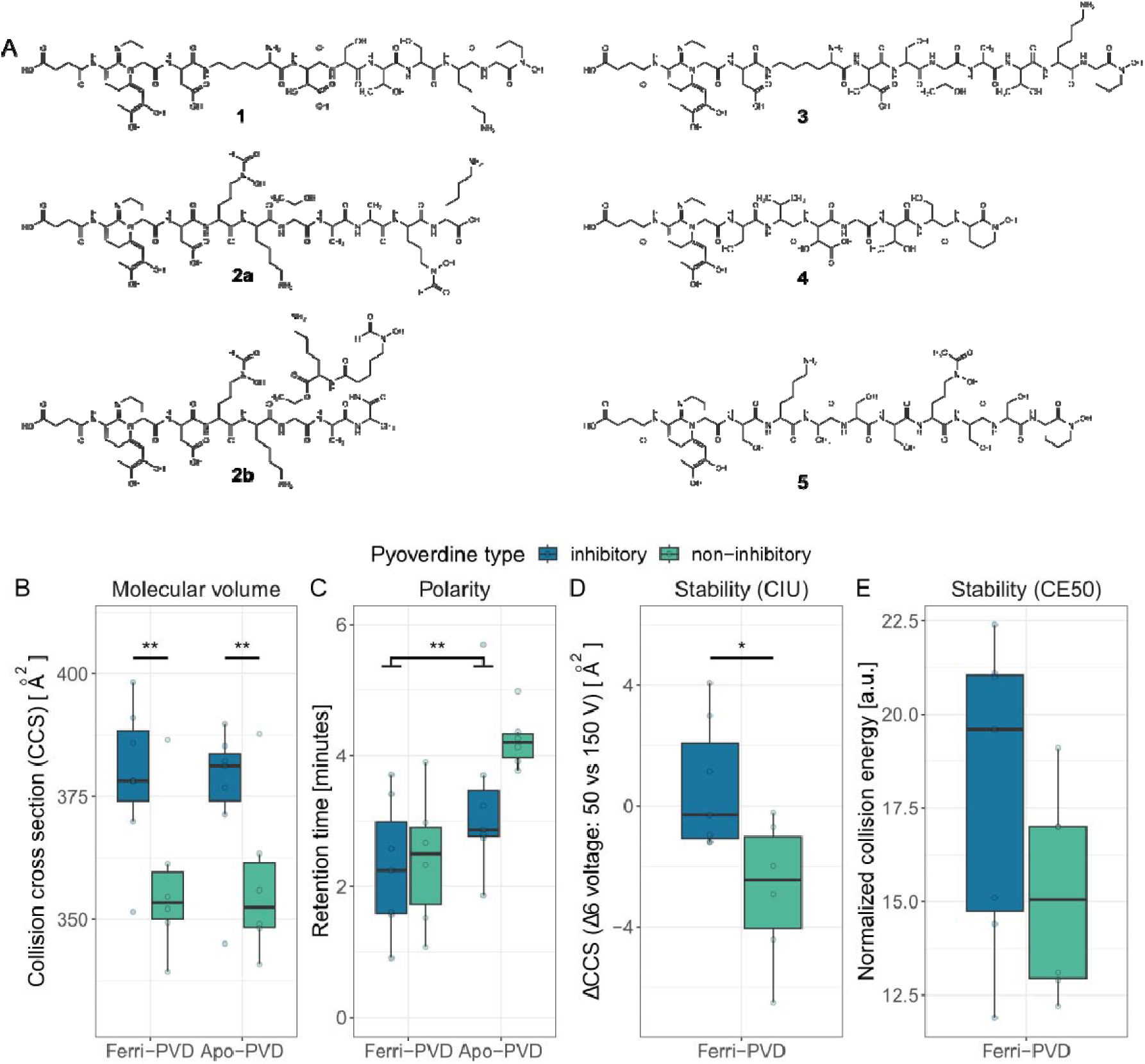
Chemical structure of growth-inhibitory pyoverdines and their properties compared to non-inhibitory pyoverdines. (A) The chemical structures of the seven top candidate pyoverdines were elucidated using UHPLC-HR-MS/MS and revealed 5 unique pyoverdine structures (**1**-**5**) differing in their peptide backbone (see^27^ for an in-depth chemical analysis). Pyoverdine **1** is a novel structure (from isolate 3A06). Pyoverdine **2** can occur in either a linear **2a** or a cyclic **2b** form (from isolate 3G07). Pyoverdine **3** was found in three different isolates, originating from the same soil sample (from isolates s3b09, s3b10 and s3b12). Pyoverdine **4** and **5** are from isolate s3c13 and s3e20, respectively. (B) The CCS values of inhibitory pyoverdines are higher than for non-inhibitory pyoverdines. (C) Molecule polarity, measured as the chromatographic retention time, is higher for ferri-pyoverdines (iron loaded) than for apo-pyoverdines (iron free). (D) Iron complex stability, assessed by the collision induced unfolding of ferri-pyoverdines, was significantly higher for inhibitory than non-inhibitory pyoverdines. (E) The normalised collision energy (CE50) necessary to fragment 50% of ferri-pyoverdines was not different between inhibitory and non-inhibitory pyoverdines. Box plots show the median and the first and third quartiles across the 7 inhibitory and 6 non-inhibitory pyoverdines. Whiskers represent the 1.5x interquartile range.

Here, we investigated whether the chemical properties of growth-inhibitory pyoverdines differ from pyoverdines that do not inhibit pathogen growth. To this end, we further elucidated the chemical structure of a set of six pyoverdines extracted from supernatants that inhibited none of the 12 pathogens, and one additional pyoverdine inhibiting the growth of 10 pathogens^27^. We found that inhibitory pyoverdines had significantly larger collision-cross sections (CCS in Å^2^) than non-inhibitory ones (F_1,22_ = 10.63, p = 0.0036, measured by ion mobility spectrometry (IMS)) (Figure 2B). As CCS values correlate with the volume of a molecule, our results suggest that inhibitory pyoverdines are larger and have more complex molecule structures than non-inhibitory pyoverdines. In contrast, there was no significant difference between the two classes of pyoverdines regarding molecule polarity (F_1,22_ = 2.00, p = 0.1711, assessed by the chromatographic retention time on a reverse phase column) (Figure 2C), but molecule polarity was significantly higher for ferri-than for apo-pyoverdines (F_1,22_ = 12.78, p = 0.0017). Finally, we assessed the stability of ferri-pyoverdines to derive a proxy for iron affinity. We first measured the propensity of ferri-pyoverdines to undergo collision-induced unfolding (CIU) by IMS measurements at different Δ6 voltages and found that inhibitory pyoverdines are significantly more stable than non-inhibitory pyoverdines (F_1,11_ = 7.55, p = 0.019) (Figure 2D). The same trend was observed for our second measure of stability (CE50, normalised collision energy necessary to fragment 50% of ferri-pyoverdines), but in this case, the difference between the two pyoverdine classes was not significant (F_1,11_ = 1.87, p = 0.198) (Figure 2E). Taken together, the chemical analyses suggest that inhibitory pyoverdines are larger and have a higher affinity for iron than non-inhibitory pyoverdines.

### Pyoverdines inhibit human pathogens in a concentration-dependent manner

We purified three out of the five most potent pyoverdines and tested their efficacy against the four human pathogens *A. baumannii*, *K. pneumoniae, P. aeruginosa* PAO1 and *S. aureus* JE2. For these experiments, we decided to include the most common (s3b09), the novel (3A06), and the cyclical-linear (3G07) pyoverdines. We crude-purified all three pyoverdines using established protocols^16,35^ and further purified the most common pyoverdine (s3b09) using preparative and analytical HPLC. During HPLC purification, we observed that ferribactin (the final pyoverdine precursor) was also highly abundant in the supernatant^36,37^, presumably released by cells during the centrifugation process. This precursor is typically not secreted but matures into pyoverdine in the periplasma^38^. Due to its compromised iron-binding capacity, we used it as a negative control for which no activity against pathogens is expected^37^. As positive treatment control, we used the antibiotic ciprofloxacin. We subjected the four pathogens to the three pyoverdine variants, ferribactin, and ciprofloxacin across a concentration gradient and measured their effects on pathogen growth (Figure 3, Figure S3).

**Figure 3.**
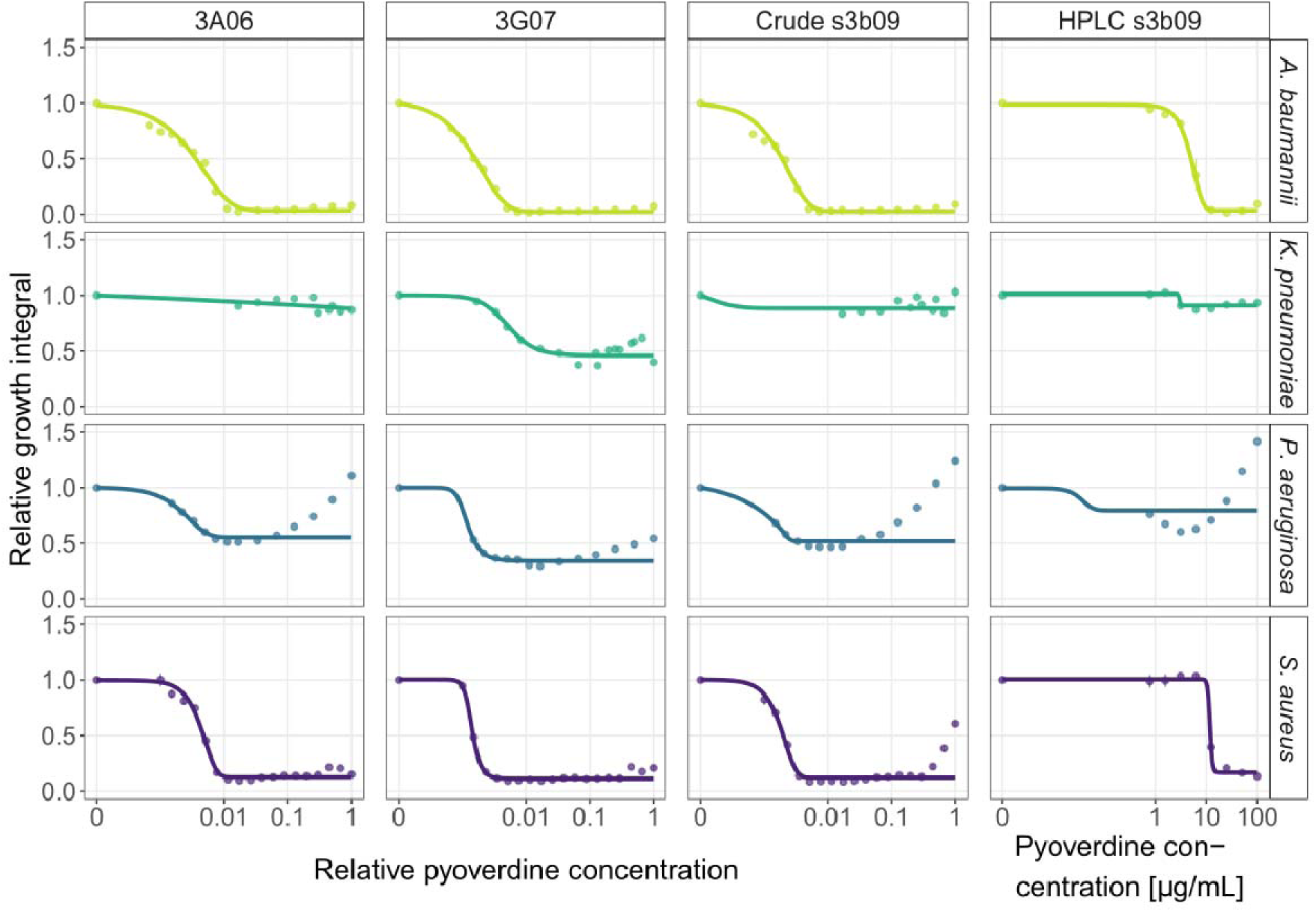
Pyoverdine dose-response curves for *A. baumannii*, *K. pneumoniae*, *P. aeruginosa* and *S. aureus*. We exposed the four human opportunistic pathogens to three pyoverdines (3A06, 3G07, s3b09) that were among the most potent ones. We used crude-purified extracts of all three pyoverdines and a HPLC-purified variant for pyoverdine s3b09. The absolute concentrations of the crude-purified extracts are unknown and therefore expressed relative to the weighed amount of 6 mg. The absolute concentration of the HPLC-purified variant is given in µg/mL. Growth values are scaled relative to the untreated control in CAA medium. Dots and error bars show mean values and standard errors, respectively, across a minimum of three replicates per concentration. Dose-response curves were fitted using 4- or 5-parameter logistic regressions.

For *A. baumannii* and *S. aureus,* we observed classical dose-response curves in all pyoverdine assays, characterized by a decrease in bacterial growth at low and intermediate concentrations, followed by a complete halt of bacterial growth at higher pyoverdine concentrations. There was a good qualitative match between the dose-response curves obtained with the HPLC vs. the crude-purified pyoverdines. Note that pyoverdine concentration is unknown for the crude extracts and is thus expressed as relative concentrations. In contrast, the pyoverdine concentration is absolute for the HPLC-purified version, allowing the determination of an IC50 value for pyoverdine s3b09, which is 5.037 µg/mL and 12.163 µg/mL for *A. baumannii* and *S. aureus*, respectively.

For *K. pneumoniae*, the pyoverdines were less effective in reducing pathogen growth. The dose-response curves for the pyoverdines s3b09 (HPLC and crude) and 3A06 showed similar trajectories and the model fits showed a growth reduction between 9-12%. The strongest inhibitory effects were found for pyoverdine 3G07 with an estimated 54% growth reduction. One reason why pyoverdines might be less effective against *K. pneumoniae* is that this species produces enterobactin, a siderophore that has higher iron affinity than pyoverdine^39^.

For *P. aeruginosa*, we observed that all pyoverdines were growth inhibitory at intermediate concentrations. The initial inhibition converted into growth promotions at high pyoverdine concentrations in three out of four cases. The only exception was pyoverdine 3G07, which yielded consistent growth inhibitions of about 66%. One reason for these non-standard dose-response curves could be that *P. aeruginosa* possesses two receptors (FpvA and FpvB) for pyoverdine uptake^40^. It might thus capitalize on the supplemented pyoverdines, especially at high concentrations via the up-regulation of the heterologous promiscuous FpvB receptor^41,42^. Our findings for *P. aeruginosa* are not unexpected and show that pyoverdine treatment should not be applied against this pathogen.

Taken together, we found that all pyoverdines are highly potent against *A. baumannii* and *S. aureus,* and one pyoverdine (3G07) is potent against *K. pneumoniae*. Especially in the case of *A. baumannii* and *S. aureus,* the pyoverdine dose-response curves follow the standard trajectory observed for antibiotics (Figure S3 for ciprofloxacin). In support of our hypothesis that the inhibitory effects of pyoverdines operate via withholding iron from pathogens, we found no growth inhibition when the pathogens were exposed to purified ferribactin (Figure S3), the non-iron chelating precursor of pyoverdine.

To further validate that the antibacterial effect of pyoverdine is due to iron sequestration, we saturated pyoverdine s3b09 with varying concentrations of iron and repeated the dose-response experiment with *A. baumannii* (Figure S4). Here, we expect pyoverdine potency to decrease because iron is no longer a growth limiting factor. Indeed, we observed that the inhibition of *A. baumannii* was greatly reduced with intermediate levels of iron supplementation (40 µm) and completely stalled with high levels of iron supplementation (200 µM) (Figure S4).

### Pyoverdines have low toxicity for mammalian cell lines, erythrocytes, and hosts

We investigated the potential toxicity of crude-purified pyoverdines against mouse neuroblastoma-spinal cord (NSC-34) hybrid cells and human embryonic kidney 293 (HEK-293) cells. NSC-34 are motor neuron-like cells, which do not divide in their differentiated state, thus mimicking established tissue. For these cells, we observed no adverse effects on cell viability at low and intermediate pyoverdine concentrations (Figure 4A), representing the doses that strongly inhibited the pathogens (Figure 3). A moderate decrease of cell viability was only observed at the highest pyoverdine concentrations (Figure 4A). Conversely, HEK-293 cells divide rapidly and inform us on whether pyoverdines interfere with cell proliferation. We indeed noticed a more pronounced effect on cell viability already at intermediate pyoverdine concentrations (Figure 4A), which suggests that pyoverdine chelates the iron required for cell proliferation.

**Figure 4.**
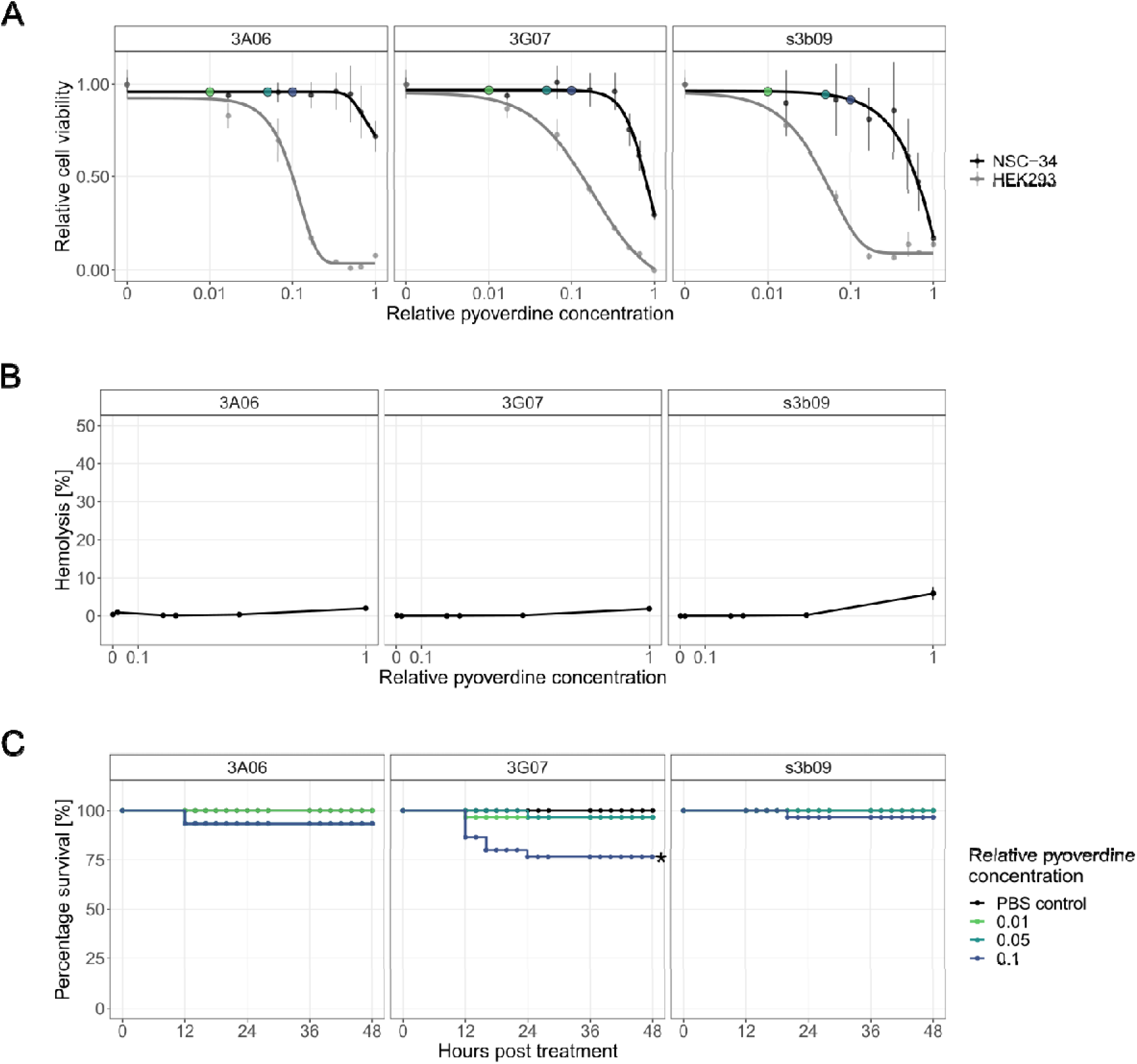
Toxicity assays for pyoverdines from environmental *Pseudomonas* spp. against human cell lines, sheep erythrocytes and the host larvae of *G. mellonella*. (A) We exposed mouse neuroblastoma-spinal cord (NSC-34) and human embryonic kidney 293 (HEK-293) cells to three crude-purified pyoverdines (3A06, 3G07, s3b09) that were among the most potent ones to inhibit bacterial growth. An MTT assay was used to assess the metabolic activity of cells as an indicator of cell viability and proliferation. Cell viability data are scaled relative to the pyoverdine-free treatment, whereby dots and error bars show means and standard error across three replicates, respectively. The absolute concentrations of the crude-purified pyoverdines are unknown and concentrations are therefore expressed relative to the highest one used. Coloured dots indicate pyoverdine dosages used for the *in vivo* experiments (pyoverdines/10 µL). Dose-response curves were fitted using 5-parameter logistic regressions. (B) We evaluated the haemolytic activity of the pyoverdines by adding them to sheep erythrocytes along a concentration gradient (range 0.002 – 1). Triton X-100 and PBS served as positive and negative control, respectively. Haemolytic activity is scaled relative to the positive control, dots and error bars show means and standard error across 6 replicates from two independent experiments, respectively. (C) To assess the toxic effects of pyoverdine on the host, we injected pyoverdines (three relative concentrations, 0.01, 0.05, 0.1) into larvae 4 hours after a mock infection with PBS. The percentage of larval survival was tracked over 48 hours post-treatment. Data stem from three independent experiments with each 10 larvae per infection and treatment.

To assess the haemolytic activity of pyoverdines, we exposed sheep erythrocytes to a concentration gradient of the crude-purified pyoverdines. We found that haemolysis was extremely low (<1%) relative to the positive control Triton X-100 surfactant for pyoverdine concentrations up to 0.5 (Figure 4B) and remained low even for the highest concentration (haemolysis rates for pyoverdines 3A06, 3G07 and s3b09 were 2.01%, 1.89% and 5.92%, respectively). This result shows that pyoverdines are unable to retrieve iron from haemoglobin.

Finally, we injected pyoverdine into the larvae of *G. mellonella* and followed host survival over time. We selected three pyoverdine concentrations (0.01, 0.05, and 0.1), which we later used for the infection experiments, and which were minimally toxic to cell lines (coloured dots in Figure 4A). We found that pyoverdine injections did not significantly reduce host survival at low and intermediate concentrations (Figure 4C and Table S1). In contrast, we observed a tendency towards higher larval mortality levels with the highest pyoverdine concentration administered, especially for 3G07, suggesting that pyoverdines can be mildly toxic for the host at high concentrations.

### Pyoverdine treatment increases the survival of infected hosts

We then investigated whether pyoverdines can be used to treat infections in *G. mellonella*. We used *A. baumannii*, *K. pneumoniae,* and *P. aeruginosa* PAO1 as pathogens. We did not include *S. aureus* for these experiments as it is literally avirulent for *G. mellonella* given the infection loads and time frames used^43^. We first determined the bacterial infection load needed to reduce larval survival by at least 50% (Figure S5) over 48 hours. We found this to be the case with 1.8*10^5^ CFU/larvae of *A. baumannii* and 8.9*10^5^ CFU/larvae of *K. pneumoniae*. For *P. aeruginosa*, all larvae died within 48 hours even with the lowest infection dose and we decided to use 56 CFU/larvae. We infected the larvae with one of the three pathogens, and subsequently treated them with either pyoverdine (relative concentrations 0.01, 0.05 and 0.1) or a PBS control four hours post infection, followed by monitoring larval survival over time (Figure 5 and Figure S6).

**Figure 5.**
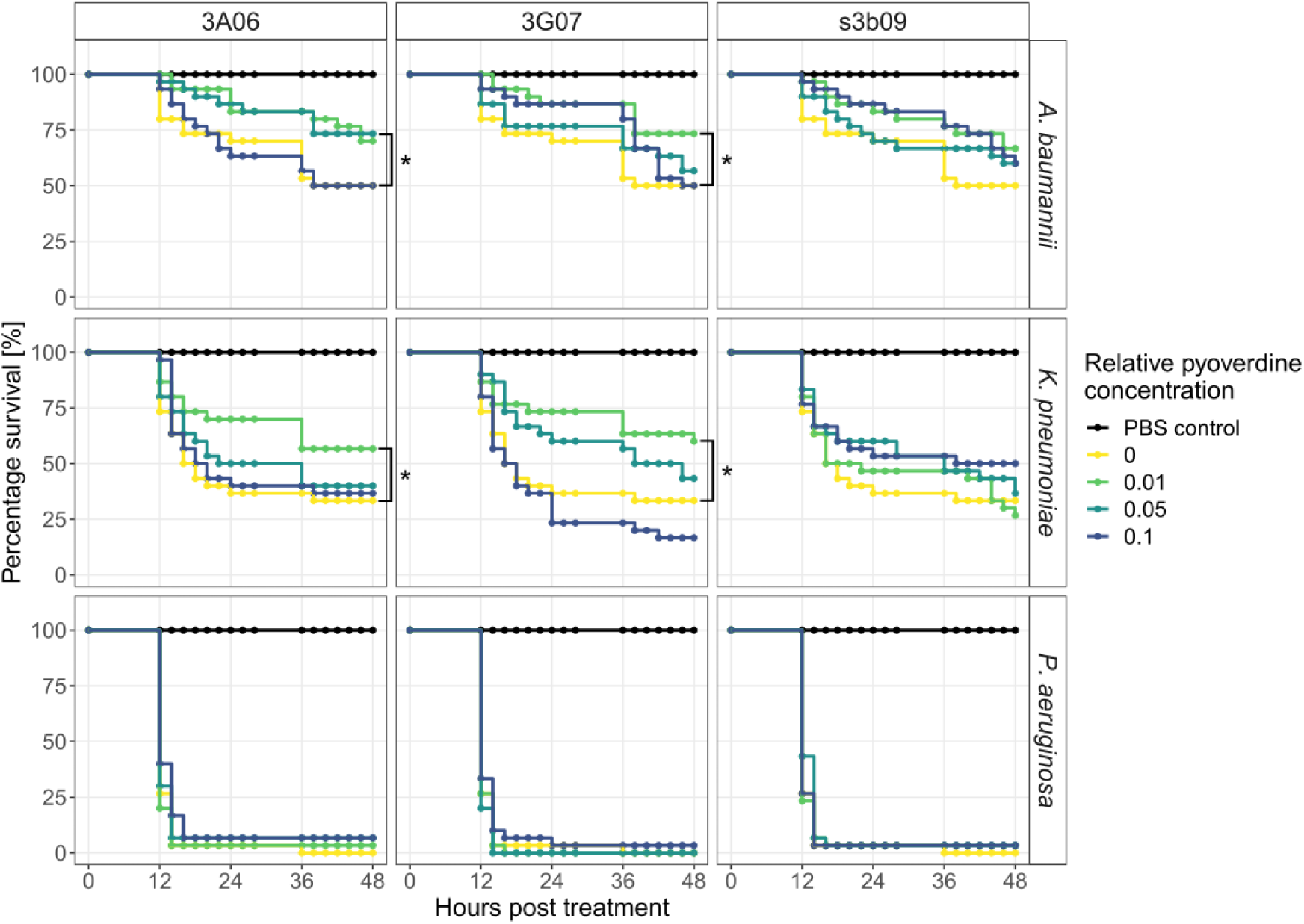
Pyoverdine treatments significantly increase survival of the host *G. mellonella* when infected with *A. baumannii* and *K. pneumoniae*. Larvae of the greater wax moth were first infected with either *A. baumannii*, *K. pneumoniae*, or *P. aeruginosa* and then treated with one of three pyoverdines (3A06, 3G07, or s3b09) at four relative pyoverdine concentrations (0, 0.01, 0.05, 0.1). All panels show the larvae survival over 48 hours post-treatment. Data stem from three independent experiments with each 10 larvae per infection and treatment. Asterisks indicate significant differences in larval survival between treated and untreated infections based on Cox proportional hazard regressions (p < 0.05).

We found that the two pyoverdines 3A06 and 3G07 were effective treatments against *A. baumannii* and *K. pneumoniae* and significantly decreased the risk of death compared to untreated infections (Figure 5, Table S2). Specifically, *A. baumannii* treated with 3A06 at a concentration of 0.05 and 3G07 at a concentration of 0.01 reduced the risk of death (hazard) by 57.9% (z = -1.973, P = 0.0485) and 59.4% (z = -2.057, p = 0.0397), respectively. For s3b09, the same tendency was observed (z = -1.496, p = 0.135) albeit not significant. For *K. pneumoniae*, the pyoverdine treatments 3A06 and 3G07 (concentration of 0.01) reduced the risk of death by 51.5% (z = -2.027, p = 0.0426) and 57.6% (z = -2.341, p = 0.0192), respectively. These results suggest that low and intermediate doses of pyoverdine are more effective than the highest concentration. This notion was further supported when analysing the time-to-death of larvae (Figure S6). We found hump-shaped relationships between pyoverdine concentrations and time-to-death in four out of six cases with two of them being significant (quadratic regression models for *K. pneumoniae* treated with pyoverdine 3G07: linear term: F_1,71_ = 0.88, p = 0.3506, quadratic term: F_1,71_ = 6.11, p = 0.0159 and s3b09: linear term: F_1,73_ = 0.31, p = 0.5806, quadratic term: F_1,73_ = 6.16, p = 0.0153). In contrast, infections with *P. aeruginosa* led to near 100% larval mortality regardless of pyoverdine treatment and dosage so that no significant treatment effect arose (Figure 5) and no effect on the time-to-death of larvae could be observed (Figure S6).

Altogether, our infection experiments show that pyoverdines are effective against the moderately virulent pathogens *A. baumannii* and *K. pneumoniae* by decreasing the risk of death by more than 50%, while they are ineffective against the fast-killing highly virulent *P. aeruginosa*. Moreover, intermediate pyoverdine concentration are most effective, reinforcing our observation that pyoverdine can exhibit some toxicity at higher concentrations (Figure 4C).

### Pathogens show low levels of resistance evolution against pyoverdine treatment

We assessed the ability of *A. baumannii*, *K. pneumoniae*, *P. aeruginosa* and *S. aureus* to evolve resistance against the pyoverdine treatment by exposing all pathogens to both single pyoverdine treatments (3A06, 3G07, and s3b09) and combination treatments (double and triple combinations) over 16 days of experimental evolution. One exception was *K. pneumoniae*, which was only exposed to 3G07 as only this pyoverdine was inhibitory. We had six independent lineages (populations) per pathogen and treatment combination (see Table S2 for treatment concentrations). We further subjected all pathogens to the antibiotic ciprofloxacin as positive control in which resistance evolution is expected. Additionally, we let the pathogens grow in untreated growth medium to control for adaptation to the growth medium. Overall, we had 180 lineages that were transferred daily to fresh medium.

After experimental evolution, we observed that all four pathogens grew significantly better under ciprofloxacin treatment than the ancestor and beyond the level observed for growth medium adaptation (Table S3; Figure 6A and S9). This provides strong evidence for pervasive resistance evolution against this conventional antibiotic. In contrast, we found that none of the four pathogens treated with pyoverdines experienced a significant increase in growth relative to the observed level of growth medium adaptation (Table S3, Figure 6A), suggesting low potentials of resistance evolution at the population level.

**Figure 6.**
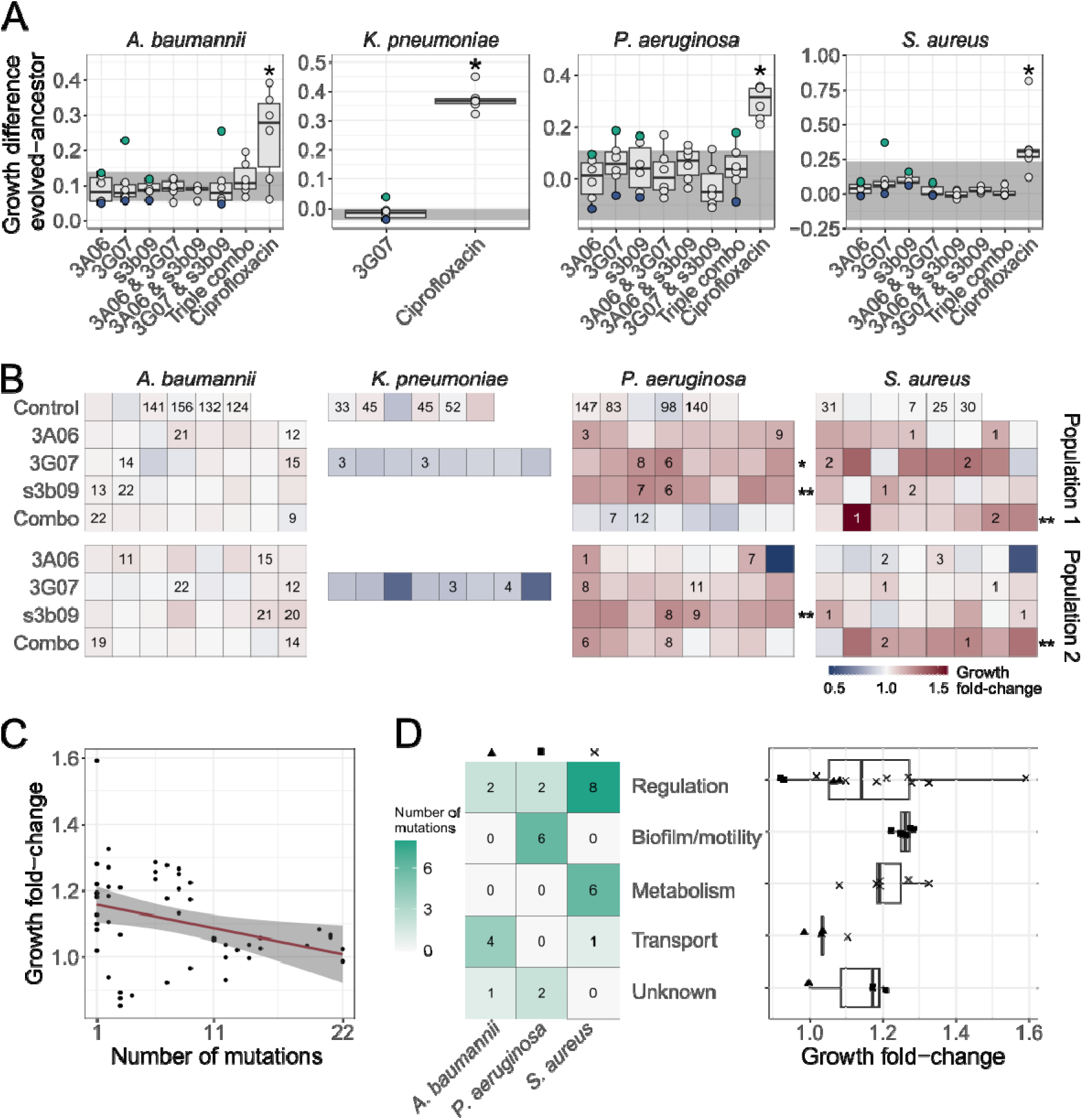
Phenotypic and genotypic analysis of experimentally evolved pathogens reveal weak levels of resistance evolution against pyoverdine treatment. (A) We exposed evolved and ancestral pathogen populations to the treatment in which they evolved in and quantified their growth (area under the curve). Growth values were scaled relative to the ancestor in untreated medium, and the panels show the scaled growth differences between evolved and ancestral populations. The shaded areas show the scaled growth difference between the ancestor and the populations evolved in untreated medium and is representative of medium adaptation. The blue and green dots represent the pathogen populations with the lowest (population 1) and highest (population 2) scaled growth difference, respectively, which were subsequently used to pick clones. The dots show mean values across two independent replicates and asterisks show significant growth increases relative to the medium-adapted control. Box plots show the median and the first and third quartiles across the 6 independently evolved populations. Whiskers represent the 1.5x interquartile range. (B) We repeated the above growth assays with 208 individual clones evolved under the pyoverdine treatments (picked from population 1 and population 2) and 24 populations evolved in growth medium alone (control). Each square represents a clone or a control population, and the number indicates the number of mutations identified based on whole-genome sequencing. The heatmap shows the fold-change in growth relative to the evolved control populations. Asterisks depict significant fold-increases in growth compared to control populations. (C) The relationship between the number of mutations and fold-change in growth across all sequenced clones (n = 52) The red line and shaded area are the regression line and 95% confidence interval, respectively. (D) Heatmap showing the number of mutations per pathogen and per functional gene categories across clones together with the respective fold-change in growth (n = 32). For this analysis, we excluded intergenic mutations and deletions larger than 600 bp. Triangle, square and cross depict growth fold-change of *A. baumannii*, *P. aeruginosa* and *S. aureus*, respectively.

However, population screens can hide the presence of resistant clones within populations. We thus picked eight random clones from two populations for each of the pyoverdine single treatments and one combo treatment for each pathogen. With a total of 208 clones, we repeated the above growth assay and expressed clonal growth as fold-change relative to the growth of populations evolved in the growth medium alone (Figure 6B and S10). For *A. baumannii* and *K. pneumoniae*, the average growth fold-change across clones was 1.02 ± 0.004 and 0.84 ± 0.02, respectively, and there was no sign of resistant clones being present in populations. This was different for *P. aeruginosa* and *S. aureus*, for which we observed high variation in growth fold-changes across clones, with many clones growing moderately better than the growth medium controls. The growth increase across clones was 1.11 ± 0.02 and 1.10 ± 0.02, and significant in three and two population of *P. aeruginosa* and *S. aureus*, respectively. Thus, the clonal screen indicates some level of adaptation to pyoverdine treatment for these two pathogens.

### Mutational and functional patterns in response to pyoverdine treatment are species specific

To obtain insights into the genetic basis of putative resistance evolution, we sequenced the whole genomes of 52 evolved clones (16 per pathogen, except 4 for *K. pneumoniae*), 16 medium-evolved control populations and the 4 ancestors. We first mapped reads to the reference genomes and then to our ancestors. Among the clones evolved under pyoverdine treatment, we identified 97 synonymous, 96 nonsynonymous, 59 intergenic and 9 nonsense single-nucleotide polymorphisms (SNPs). Additionally, there were 48 insertions, 59 deletions and 45 substitutions (Figure 6B). We observed that growth peaked in clones with a single mutation and decreased in clones with multiple mutations (Figure 6C, linear fit: F1,50 = 6.57, p = 0.0135), suggesting positive selection for a few specific advantageous mutations.

Next, we excluded all mutations that were present at least twice in the no-treatment controls, and all intergenic mutations. We identified seven mutations in *A. baumannii*, one in *K. pneumoniae*, ten in *P. aeruginosa*, and 20 in *S. aureus* that exclusively arose under pyoverdine treatment and could thus be associated with resistance evolution. We then classified the SNPs and small deletions (<600 bp) into functional categories (Figure 6D, Table S4). We found that mutations in genes related to regulation and metabolism were consistently associated with increased growth in *S. aureus*, whereas mutations in genes related to biofilm and motility were associated with increased growth in *P. aeruginosa* (Figure 6D).

Furthermore, we identified six large deletions (≥600 bp; Table S5). One *K. pneumoniae* clone lost a ∼42 kb plasmid encoding various functional proteins, such as methyltransferases and toxin-antitoxin systems^44^. Four *S. aureus* clones from two independent populations showed parallel loss of the ∼14 kb pathogenicity island SaPI5, containing the virulence genes *sek* and *seq*^45,46^. Additionally, one *S. aureus* clone exhibited a ∼55 kb deletion containing the staphylococcal cassette chromosome mec type IV (SCCmecIV), which harbours the *mecA* gene responsible for β-lactam antibiotic resistance, and the arginine catabolic mobile element (ACME)^45^.

Taken together, we found that the observed weak resistance evolution against pyoverdine in *P. aeruginosa* and *S. aureus* was not linked to mechanisms directly enhancing the pathogen’s iron acquisition but with mutations in global regulators, metabolism, biofilm and motility genes. In *A. baumanii* and *K. pneumoniae*, no evidence for resistance evolution was observed. Moreover, certain pathogens lost mobile genetic elements containing virulence and antibiotic resistance genes, suggesting that pyoverdine treatment could selects for less virulent and antibiotic sensitive strains.

## Discussion

We investigated whether pyoverdines produced by environmental *Pseudomonas* spp. have antibacterial activities against human opportunistic pathogens through iron sequestration and withholding this trace element from pathogens. Our screen involving 320 environmental isolates and 12 pathogen strains revealed five top pyoverdine candidates with broad-spectrum activity. These candidates could be distinguished from non-inhibitory pyoverdines by their high CCS values (standing for larger and more complex molecules) and their higher iron complexation stability. Experiments with crude- and HPLC-purified pyoverdines showed that pyoverdines completely stall the growth of *A. baumannii* and *S. aureus*, while showing intermediate activity against *K. pneumoniae* and *P. aeruginosa*. When administered as treatment to infected *G. mellonella* larvae, we observed significantly increased host survival rates in infections with *A. baumannii* and *K. pneumoniae*, demonstrating that pyoverdine exhibits antibacterial activity in hosts. Furthermore, we found low toxicity of pyoverdines at effective concentrations and observed low potentials for resistance evolution. Overall, our results reveal pyoverdines from non-pathogenic *Pseudomonas* spp. as potent antibacterials against several human opportunistic pathogens.

Our novel treatment approach assumes that pyoverdines sequester iron and thereby induce iron starvation and growth arrest in pathogens. Several of our findings support this view. First, the growth-inhibiting effect of our top pyoverdine candidates exclusively occurred under iron-limited conditions, while pathogen growth was restored under iron-replete conditions (Figure 1, Figure S4). Second, non-inhibitory ferri-pyoverdines were more prone towards collision induced unfolding than inhibitory ferric-pyoverdines (Figure 2D), suggesting that inhibitory pyoverdines have a higher iron binding affinity. Third, the pyoverdine precursor ferribactin did not inhibit pathogen growth (Figure S3). The two molecules are identical apart from one ring in the chromophore core that is not cyclized and not oxidized in ferribactin, which compromises the binding of iron. These findings suggest that pyoverdine does not target the pathogens directly, but rather indirectly via the induction of iron starvation.

We observed that pyoverdines showed pathogen-specific efficacies in curbing bacterial growth (Figure 3). We propose that differences can be explained by the various strategies of these pathogens to cope with iron stress. *A. baumannii* and *S. aureus* produce siderophores that have a much simpler chemical structures than pyoverdines and are expected to bind iron with lower affinity (*A. baumannii*: acinetobactin (association constant: 10^26^ M^-1^), fimsbactin (10^27^ M^-1^) and baumannoferrin (unknown)^47–49^, *S. aureus*: staphyloferrin A+B (unknown)^50^) than pyoverdine (10^32^ M^-1^)^28^. Hence, the growth suppression of *A. baumannii* and *S. aureus* most likely occurs because their siderophores are too weak to successfully compete with pyoverdine. The situation is different for *K. pneumoniae*, which produces enterobactin, the siderophore with the highest known iron affinity (10^52^ M^-1^)^39^. Consistent with the concept of iron competition, we observed that pyoverdine treatment is less effective against *K. pneumoniae*, and was only potent for one out of three pyoverdines (3G07, Figure. 3). Finally, pyoverdine treatment is not expected to work well against *P. aeruginosa*, which itself produces a version of this siderophore. The two competing pyoverdines likely have similar iron affinities and the presence of pyoverdine receptors in the pathogen might foster the uptake of the supplemented pyoverdine. Our results indeed support the view that pyoverdine treatment is not effective against *P. aeruginosa*. Taken together, we predict that pyoverdine treatment could be highly potent against certain pathogens like *A. baumannii* and *S. aureus* that have less efficient siderophores and are unable to use the pyoverdine as iron source.

For every new treatment, it is important to consider whether it has unintentional consequences for the targeted pathogen. Because our approach does not kill pathogens directly, there are multiple ways of how pathogens may react to pyoverdine treatment. For example, pathogens could increase the production of their own siderophores in response to the severe iron limitation induced by pyoverdines. Alternatively, siderophores can serve as signalling molecules in certain species like *P. aeruginosa*, regulating the expression of additional virulence factors like proteases and toxins^51^. While pyoverdine treatment may affect such signalling cascades in other pathogens as well, and thus potentially increase virulence factor production, we observed the opposite in *S. aureus*, which showed a loss of virulence genes. Similarly, iron limitation can increase^52^ or decrease^53–55^ the formation of biofilms, depending on the species or even strains^56,57^ and it is well possible that pyoverdine treatment thus affects the propensity of pathogens to form biofilms. Finally, pyoverdines may have differential effects on the various members of the microbiota or the different pathogens in polymicrobial infections and thereby induces shifts in species composition due to the unequal suppression of community members. In complex communities, the inhibitory effect of pyoverdine on the pathogen may get diluted as the burden of increased iron limitation is shared across community members. All these considerations show that it will be crucial to assess the consequences of pyoverdine treatments in more complex settings including hosts.

Not only bacteria but also host cells need iron. Consequently, pyoverdine treatment could negatively affect host iron homeostasis. Our toxicity assays reveal that there is a range of pyoverdine concentrations for which only mild adverse effects against cell lines and *G. mellonella* larvae are observed and no haemolysis occurs. Especially the finding that pyoverdines have no negative effects on the survival of non-replicating NSC-34 cells is encouraging, as this cell line is representative of intact host tissue, in which iron is bound to strong chelators such as transferrin, lactoferrin or ferritin^58,59^. Additionally, recent research has demonstrated that purified pyoverdine remains non-toxic even after 72 hours of treatment at concentrations up to 200 μM^60^. The result that pyoverdines are unable to obtain iron from haemoglobin is encouraging, too. Previous work showed that the pyoverdine of *P. aeruginosa* can retrieve iron from human transferrin when working in concert with proteases (proteolytically degrading transferrin)^61^ or phenazines (spontaneously reducing ferric to ferrous iron)^62^. However, these *P. aeruginosa* specific mechanisms require the presence of this pathogen^63^, which is not the case when purified pyoverdines are used as a treatment against other pathogens like *A. baumannii*. Nonetheless, we found that pyoverdines can have certain adverse effects, especially for the rapidly proliferating HEK293 cells (Figure 4). These cells represent a regenerating tissue and thus have high iron demands. Similarly, we observed mild adverse effects for some pyoverdines (3G07) when administered at high concentrations to *G. mellonella* larvae (Figure 5). These findings corroborate the results from previous studies on *Caenorhabditis elegans*, where pyoverdines were found to interfere with host iron homeostasis^64,65^. These considerations highlight that finding the right pyoverdine concentration will be key to meet the fine line between maximizing pathogen inhibition and minimizing tissue damage. By contrast, the adverse host effects of pyoverdines might come with additional opportunities, for instance in cancer therapy, where siderophores reduce iron levels in tumours and slow tumour progression^66^.

We propose that pyoverdine treatment could synergistically interact with the host innate immune system. Mammalian neutrophils produce siderocalin to sequester bacterial siderophore-iron complexes, rendering them non-functional^58^. While this strategy is efficient against many siderophores, some bacteria have evolved so-called stealth siderophores that no longer bind to siderocalin. Pyoverdines are such stealth siderophores^67^. Regarding pyoverdine treatment, this means that pathogens with regular siderophores will hit a double wall: their siderophores are immobilized by siderocalins, while pyoverdines lock away the remaining iron.

Another promising finding of our work is that the potential for resistance evolution against pyoverdines seems low. Conceptually, one reason for low resistance evolution could be that pyoverdines, unlike many antibiotics, are not internalized into bacterial cells. Thus, classic resistance mechanisms such as reduced drug influx, increased drug efflux, and intra-cellular target modification cannot operate^68^. We initially expected that resistance evolution would involve mechanisms that improve the pathogen’s iron acquisition. This could include the up-regulation of the pathogen’s own siderophores or switching from siderophore-based ferric iron acquisition to the use of reductases that foster ferrous iron uptake^68^. However, our sequencing analysis did not reveal mutations in genes associated with iron uptake systems. One explanation is that such mutations (albeit beneficial) might not reach high frequencies in populations because increased siderophore production and secretion would increase the iron acquisition of resistant and non-resistant cells alike^68^. We instead observed an overrepresentation of mutations in regulatory, biofilm and motility, and metabolic genes. While we do not know the phenotypes of these mutations, one possibility is that increased biofilm formation could offer some protection against pyoverdine treatment, for example by restricting pyoverdine from chelating free iron. Important to note is also that patterns of resistance evolution may differ between *in vitro* and *in vivo* settings^4,69^. For example, our experimental evolution setup did not allow for horizontal transfer of siderophore receptors, a scenario that might occur in the context of polymicrobial infections and confer resistance to pyoverdine therapy.

In conclusion, our treatment approach using pyoverdine is based on the principle of paying like with like. Pathogens secrete siderophores to scavenge iron for their growth. We add a strong heterologous siderophore as treatment to withhold iron from pathogens to curb their growth. The concept of targeting bacterial infections with both natural and synthetic iron-chelators has been highlighted in the literature for quite some time^70–74^. For instance, the siderophore desferoxamine (produced by *Streptomyces* species) is used in clinical settings to treat iron overload^75^. However, its efficacy as antibacterial is limited because of its low iron affinity (10^27^ M^-1^, compared to pyoverdine and enterobactin)^76^ and several pathogens have receptors for deferoxamine uptake and can thus use it as an iron source^77^. Synthetic iron chelators like deferiprone and deferasirox have been proposed as alternatives to overcome these limitations. However, the minimum inhibitory concentration (MIC) of deferiprone needed to inhibit *P. aeruginosa* was found to be up to 36.7 times higher than the plasma levels in human therapy, raising concerns about its cytotoxicity and clinical viability for treating bacterial infections^78,79^. Furthermore, at sub-MIC levels, deferiprone only partially suppressed the growth of *P. aeruginosa* and *A. baumannii,* indicating insufficient iron sequestration from these pathogens^79,80^. Our approach offers a distinct advantage by utilizing naturally evolved iron chelators from non-pathogenic species, which possess unique chemical structures and exceptionally high iron affinities. These properties reduce the likelihood of their exploitation as iron sources by pathogens and provide a competitive advantage over siderophores with lower iron affinities. Combined with low toxicity to hosts at therapeutically effective concentrations and minimal potential for resistance evolution, these natural chelators hold promise as a new class of effective antibacterials, either as standalone treatments or in combination with conventional antibiotics.

## Material and methods

### Bacterial strains

We used a strain collection of 320 natural isolates, originating from soil and pond habitats. The isolates originate from 16 independent soil and water samples (20 isolates per sample) collected on the Campus Irchel Park of the University of Zurich. In previous work, we have extensively studied and characterized all isolates^16,29–31^. Details on the isolates can be found in the Supporting Information.

For the supernatant screening assay, we used a collection of 12 strains of opportunistic human pathogens (Table 1), including *A. johnsonii*, *A. junii*, *C. sakazakii*, *K. michiganensis* (belonging to the *K. oxytoca* complex)^81^ and *Shigella* sp. (all provided by the laboratory of Leo Eberl, University of Zurich), and *B. cenocepacia* strains H111 and K56-2, *P. aeruginosa* strains PA14 and PAO1, *E. coli* K12 and *S. aureus* strains Cowan and JE2 from our own strain collection. For dose-response and infection experiments, we used *A. baumannii* (DSM 30007) and *K. pneumoniae* (DSM 30104), both purchased from the German Collection of Microorganisms and Cell Cultures GmbH, in addition to *P. aeruginosa* PAO1 and *S. aureus* JE2.

### Culturing conditions

Pathogen overnight cultures were grown in either 8 mL tryptic soy broth (TSB; for the two *S. aureus* strains) or 8 mL lysogeny broth (LB; all other pathogens) in 50 mL tubes at 37 °C and 220 rpm agitation. Following two washing steps with 0.8% NaCl, we adjusted the overnight cultures to an optical density at 600 nm (OD_600_) of 1. For overnights of environmental isolates, we followed the same procedure when working with low sample sizes, with the only difference being that culturing occurred at 28 °C. For the large-scale screening experiments, we grew environmental isolates in 200 µL LB in 96-well plates shaken at 170 rpm. Cultures from plates were then directly used for experiments. For all main experiments (screen for bioactive pyoverdines and dose-response curves), we used casamino acid (CAA) medium (1% casamino acids, 5 mM K_2_HPO_4_ * 3H_2_O,1 mM MgSO_4_*7 H_2_O, 25 mM HEPES buffer). This medium contains low levels of iron, triggering pyoverdine production^82^.

### Supernatant screening assays

We created sterile supernatants from all the 320 *Pseudomonas* isolates. Specifically, we transferred 2 µL of overnight cultures (grown in 96-well plates in LB in four-fold replication for 24 hours as described above) to new plates containing 200 µL of CAA medium supplemented with 250 µM 2,2’-bipyridyl. Cultures were incubated for 24 hours and subsequently centrifuged at 2250*g* for 10 minutes. The supernatants were transferred to 0.2 µm membrane filter plates (PALL AcroPrep Advance), centrifuged a second time, and frozen at -20 °C.

For the supernatant screening assay, the pathogens were cultured and diluted as described above and added to 140 µL CAA medium in 96-well plates. We added 60 µL of thawed *Pseudomonas* supernatants (30%) or 0.8% NaCl as control and incubated the plates at 37 °C and 170 rpm. Growth was measured as OD_600_ after 24 hours using a Tecan Infinite M-200 plate reader (Tecan Group, Männedorf, Switzerland). We then scaled the growth of pathogens in the supernatant relative to their growth in the control treatment. Relative growth values smaller and larger than one indicate growth inhibition and promotion by the supernatant, respectively.

To test whether pyoverdine is responsible for pathogen growth reduction, we repeated the screening assay but supplemented the CAA medium containing the *Pseudomonas* supernatants with 40 µM FeCl_3_. In this iron-rich condition, pyoverdine should not be able to limit iron availability and thus pathogen growth should be restored.

### Structure elucidation of pyoverdine and analyses of chemical properties

From the seven top supernatant candidates that inhibited all twelve pathogens, we extracted the pyoverdines and elucidated their structure. For this purpose, we have developed a new protocol that allows structure elucidation from low volume supernatants using UHPLC-HR-MS/MS. The methodological details are described in Rehm *et al.*^27,34^. Afterwards, we assessed whether growth inhibiting pyoverdines can be distinguished from non-inhibiting pyoverdines by their chemical properties. We focussed on polarity, CCS values, and stability upon collision induced unfolding or fragmentation. Detailed protocols and instrument settings can be found in the Supporting Information.

### Crude pyoverdine purification

To test the efficacy of the pyoverdines against the four focal human opportunistic pathogens (*A. baumannii*, *K. pneumoniae*, *P. aeruginosa* (PAO1), and *S. aureus* (JE2)), and to assess their cytotoxicity against human cell lines, we crude-purified three pyoverdines (3A06, 3G07, s3b09). The three pyoverdines were among the five most potent ones identified in our supernatant screening assay. The adapted method for pyoverdine purification from the previous studies^16,35^ are further described in the Supporting Information.

### HPLC purification of pyoverdine and ferribactin

To further validate that pyoverdines are the compounds that induce pathogen growth inhibition, we purified the most common pyoverdine (s3b09) among the most potent pyoverdines from our screens using reversed-phase high-performance liquid chromatography (HPLC). Methodological details thereof are described in the Supporting Information.

### Dose-response curves

To determine the inhibition potential of crude- and HPLC-purified pyoverdines, as well as ferribactin, we subjected *A. baumannii*. *K. pneumoniae*, *P. aeruginosa* and *S. aureus* to serially diluted extracts. For this, the extracts were dissolved in CAA medium, filter sterilized and diluted in the same medium. Bacterial overnight cultures were grown and prepared as described above and added to a total of 200 µL medium on a 96-well plate with 4 replicates per extract dilution. As positive control where growth inhibition is expected, we subjected all three pathogens to a dilution series of the antibiotic ciprofloxacin (highest concentration 4 µg/mL), in the exact same way as for the extracts. We incubated the plates at 37 °C statically in the plate reader and measured the OD_600_ every 15 minutes for 24 hours. Before each measurement, plates were shaken for 15 seconds. We repeated the experiment up to five times with different initial extract starting concentrations. This was done to ensure that the full dose response to pyoverdine was captured. Thus, sample size per individual extract concentration varied between four and 20 replicates. We subtracted the blank values and the background values caused by the treatment of the respective extract concentration from the measured growth data and calculated the integral (area under the growth curve) using the R package Growthcurver^83^. We then expressed growth relative to the control medium, where bacteria grew in CAA without extract or antibiotic addition. Finally, we plotted extract/antibiotic concentration vs. relative growth and fitted dose-response curves (see below).

### Cytotoxicity assays

To determine the cytotoxicity of pyoverdine treatments against mammalian cells, we subjected human embryonic kidney 293 (HEK-293) and mouse motor neuron-like neuroblastoma-spinal cord hybrid (NSC-34) cells to the crude extract of the three most potent pyoverdines (3A06, 3G07, s3b09). Detailed culturing conditions can be found in the Supporting Information. Briefly, approximately 10^4^ cells/well were plated on 96-well plates (Greiner Bio-One, 655090) and, after 48 hours, the culture medium was replaced and the pyoverdines were added covering a range of concentrations. After another 48 hours, the pyoverdine-containing culture medium was removed, and cells were incubated with 1 µg/mL thiazolyl blue tetrazolium bromide (Sigma, D5655) in 100 µL of fresh medium for 1.5 hours (NSC-34) or 40 minutes (HEK-293). The reaction was stopped, and formazan crystals simultaneously solubilized with the addition of 100 µl of a solution containing 10% sodium dodecyl sulfate (SDS; Sigma-Aldrich, 05030) and 0.03% HCl (Supelco, 100319). Finally, the absorbance of cell debris and other contaminants at 630 nm were subtracted from the absorbance of the solubilized formazan at 570 nm.

### Haemolysis assay

To determine the haemolytic activity of pyoverdines, we quantified sheep erythrocyte haemolysis at different pyoverdine concentrations. We first centrifuged 25 mL of fresh sheep blood (Chemie Brunschwig AG, Switzerland) in a 50 mL tube at 1500*g* for 5 minutes at 4 °C. Plasma was then gently aspirated, replaced by PBS and tubes were gently inverted to mix. This washing procedure was repeated three times. Pyoverdines were diluted in PBS to final relative concentrations of 1, 0.5, 0.25, 0.2, 0.02, and 0.002. PBS and 0.1% Triton X-100 (Sigma Aldrich, Switzerland) served as negative and positive control, respectively. 100 µL erythrocyte solution was mixed with either 100 µL pyoverdine solution or the controls in a 96-well plate in triplicates. Following an incubation at 37 °C and 120 rpm for 1 hour, the plate was centrifuged at 1036*g* for 5 minutes at 4 °C. Subsequently, 100 µL of the supernatant were transferred to a new 96-well plate. Haemoglobin release was measured at 570 nm in the plate reader and the percent haemolysis was calculated relative to the positive control (% haemolysis = (OD570 of sample – negative control) / (positive control – negative control)).

### Infection model

We tested the efficacy of pyoverdine to treat infections in larvae of the host model *G. mellonella*. Detailed information on the larvae handling and infection procedure are described in the Supporting Information. On the pathogen side, we determined the LD_50_ (lethal dose, killing 50% of the larvae) for *A. baumannii* (1.8 * 10^5^ CFU/larvae) and *K. pneumoniae* (8.9*10^5^ CFU/larvae) (Figure S5). For *P. aeruginosa*, we use 56 CFU/larvae because all doses tested killed 100% of the larvae (Figure S5). As negative control, we administered 10 µL/larvae of sterile PBS. Following infections, larvae were individually placed in wells of 24-well plates and incubated at 37 °C in the dark for 4 hours.

On the treatment side, we used pyoverdines 3A06, 3G07, and s3b09 at three concentrations that were non-toxic for the NSC-34 cells. For this, we prepared stocks of 60 mg/mL, 30 mg/mL, and 6 mg/mL of crude pyoverdines. Since the injected treatment volume was 10 µL per larvae, the final pyoverdine concentrations were 0.6 mg, 0.3 mg, and 0.06 mg per larvae, respectively. When scaling theses concentrations to the doses used for the *in vivo* dose-response curves (Figure 4), the relative pyoverdine concentrations were 0.1, 0.05 and 0.01. Pyoverdine treatments were administered 4 hours after the infection through injections (see Supporting Information). Prior to treatment, larvae were immobilized on ice for 15 minutes. We also applied treatments to larvae infected with sterile PBS to quantify potential negative effects of the treatment in the absence of an infection. Following treatment, larvae were put back to their allocated well of the 24-well plate and kept at 37 °C in the dark.

From the 12^th^ hour post infection (hpi) onwards, we regularly checked the survival of all larvae for a total of 48 hpi. Specifically, larvae were poked with a pipette tip and individuals not responding to the physical stimulation were considered dead. We conducted three independent experiments per pathogen-treatment combination with 10 larvae per experiment, resulting in a total of 30 larvae per pathogen-treatment combination. Larvae in their last instar stage were purchased from a local vendor (Bait Express GmbH, Basel, Switzerland) and different batches of larvae were used for the three independent experiments.

### Experimental evolution experiment

We first determined the IC50 concentrations of (crude) pyoverdines 3A06, 3G07, and s3b09, and the antibiotic ciprofloxacin against all four pathogens. For this, we fitted logistic regressions to the dose-response curves. We used these IC50 values as treatment concentrations during experimental evolution (Table S2). We also mixed different pyoverdines in double and triple combo treatments. In combo treatments, pyoverdine concentrations were equivalent to those of the single treatments, meaning that we divided the single treatment concentrations by two or three. The experimental evolution experiment included (i) three single pyoverdine treatments (3A06, 3G07, s3b09), (ii) three double pyoverdine treatments (3A06 and 3G07, 3A06 and s3b09, 3G07 and s3b09), (iii) one triple pyoverdine treatment (3A06, 3G07 and s3b09), and (iv) one ciprofloxacin treatment for *A. baumannii*, *P. aeruginosa* and *S. aureus*. For *K. pneumoniae*, which was only affected by one pyoverdine (3G07), the experiment consisted of one single pyoverdine treatment and one ciprofloxacin treatment.

A detailed description of the experimental evolution setup is provided in the Supporting Information. Briefly, for each pathogen-treatment combination, we included six independently replicated lineages (populations) on 96-well plates. Each plate further contained control populations that evolved in plain CAA medium. At the start, we grew the pathogens overnight, washed and diluted them as described above, and transferred 2 µL of the diluted pathogens into 200 µL medium in 96-well plates. Subsequently, the plates were incubated at 37 °C for 22 hours, shaken at 170 rpm. After incubation, we measured growth (OD_600_) of all evolving lineages with the plate reader. We then diluted the evolving cultures 1:1,000 in CAA medium and added them to a fresh treatment plate. After the transfer, the new plates were again incubated at 37 °C and 170 rpm agitation for 22 hours. We added glycerol to the old plates to a final concentration of 25% and stored them at -80 °C.

### Phenotypic screening of evolved populations

We screened evolved populations for improved growth performance relative to the ancestor under pyoverdine treatment, which we took as a proxy for resistance evolution. For this purpose, we thawed the final plates (day 16) of the experimental evolution experiment and transferred 2 µl of all populations to 200 µl of LB or TSB (for *S aureus*) to create overnight cultures in 96-well plates. We further included ancestor cultures of all four species. After incubation for 18 hours at 37 °C and 170 rpm, we diluted overnights 1:1,000 with 0.8% NaCl and subjected all evolved populations to the conditions they had evolved in. Ancestors were simultaneously exposed to the respective treatment conditions. The culturing volume and conditions matched those used during experimental evolution, but this time we incubated the plates in a plate reader, measuring OD_600_ every 15 minutes to obtain growth kinetics. The population screen was repeated twice independently. From the growth kinetics, we first calculated the area under the curve (AUC) for each growth curve. AUC values were then scaled relative to the growth of the ancestor in the untreated CAA control medium. For each evolved population, we then calculated the difference in growth in comparison to the mean growth of the ancestor subjected to the same treatment.

### Phenotypic screening of evolved clones

To screen for resistant clones within populations, we selected the pathogen populations with the lowest and highest growth difference for each single treatment and for one combination treatment and picked 8 random colonies. We transferred them to 200 µL LB or TSB (*S. aureus*) in 96-well plates and incubated plates for 18 hours at 37 °C (170 rpm) to create overnight cultures. We followed the same protocol as above for quantifying the difference in growth of evolved and ancestral clones.

### Genomic analyses of evolved populations and clones

We sequenced the genomes of the four ancestors, four control populations that had evolved in medium without pyoverdine treatment, and two random clones (out of the eight picked) per population that had received pyoverdine treatment and were used for clonal growth analysis (Figure 6B). The Supporting Information features a detailed description of the sequencing and data analysis protocols used. Variants were predicted relative to the reference genomes ATCC 19606 (*A. baumannii*), ATCC 13883 (*K. pneumonia*), ATCC 15692 (*P. aeruginosa*) and CP020619.1 (*S. aureus*) using the breseq pipeline^84^. In the pipeline, we specified that medium-adapted controls were treated as populations, whereas the treatment-adapted colonies picked for sequencing were treated as clones.

### Data analyses

All statistical analyses were performed in R 4.0.2 and RStudio version 1.3.1056. We conducted Shapiro-Wilk tests and consulted diagnostic plots to check whether our data (residuals) were normally distributed. For normally distributed data sets, we used linear models and two-sided tests for all analyses. For non-normally distributed data sets, we used non-parametric rank tests. We used ANOVA to test whether chemical properties differ between growth inhibiting and non-inhibiting pyoverdines, in their ferri-versus apo-pyoverdine state. For the *in vitro* dose-response and cytotoxicity experiments, we fitted dose-response curves with either 4- or 5-parameter logistic regressions using the nplr package^85^. Survival analyses were performed using the Cox proportional hazards regression model or the log-rank test (adjusted for multiple comparisons using the Holm method) from the survival package^86^. We used two-sided t-tests, Welch’s t-tests, or Wilcoxon tests to compare growth differences between the ancestor and the evolved populations under treatment. To compare growth differences between the ancestor and the evolved clones under treatment, we used one- and two-way ANOVAs. To construct the cladogram in Figure S2, we followed the procedure described in Kramer et al^31^ using partial *rpoD* gene sequences with lengths ≥600 bp. We used iTOL tool^87^ for the graphic presentation of the cladogram and to link phylogeny to the supernatant effects.

## Acknowledgements

We thank Richard Allen, Dominik Bär, Johanna Giger, Kevin Schiefelbein and Julien Weber for laboratory assistance, and Markus Seeger for insightful discussion. This work was supported by funding from the Swiss National Science Foundation (grant numbers: 31003A_182499 to RK, PP00P3_144862 and PP00P3_176966 to MP) and a Candoc Grant (Forschungskredit) from the University of Zurich (to MPB).

## Authors contributions

VV and RK conceived the project. VV, KR, CC and MPB performed the experiments and analysed experimental results. All authored interpreted the data. VV and RK wrote the manuscript with contributions from all authors.

## Data availability

The raw data underlying the figures are available from the Figshare depository (https://doi.org/10.6084/m9.figshare.26388565). The sequencing data for this study have been deposited in the European Nucleotide Archive (ENA) at EMBL-EBI under accession number PRJEB78468 (https://www.ebi.ac.uk/ena/browser/view/PRJEB78468).

