## Supplemental Material for "Antimicrobial activity of iron-depriving pyoverdines against human opportunistic pathogens"

**This file contains the following supporting information:**

- Supplementary Methods
- 10 supporting Figures
- 5 supporting Tables

**Supplementary Methods**

*Bacterial strains*

We used 315 isolates where sequencing of the *rpoD* housekeeping gene confirmed that they belong to the group of fluorescent pseudomonads that are non-pathogenic to humans^1^. For the remaining five isolates, the *rpoD* gene could not be amplified, yet they were still included in our study. Moreover, siderophore screens previously revealed that a large proportion of the isolates can produce and secrete pyoverdines^1^, the high-iron affinity siderophores we focus on in our study. The exact sampling and isolation procedures together with the pyoverdine production profiles are described elsewhere^1,2^. All bacterial natural isolates and pathogenic species used in our study will be available from the authors upon request.

*Structure elucidation of pyoverdine and analyses of chemical properties*

To analyse chemical properties of pyoverdines, we first purified them from the supernatants using solid-phase extraction. Next, we ran the samples through an HSS C18 column of the Vanquish UHPLC system (Thermo Fisher Scientific, Waltham, MA, USA) using formic acid (0.1%) as an eluent additive and MeCN as the organic phase and recorded the retention times (as a measure for polarity). We also measured the CCS values of pyoverdines and ferri-pyoverdines on a trapped ion mobility spectrometry (TIMS)-TOF-MS instrument (timsTOF Pro™, Bruker, Bremen, Germany) at a Δ6 voltage of 100 V. Finally, we assessed the stability of ferri-pyoverdines with two approaches. First, we investigated the proneness of ferri-pyoverdines to undergo collision induced unfolding (CIU). For this, we determined the difference in CCS values of the ferri-pyoverdines (ΔCCS) at a Δ6 voltage of 50 V (no structural unfolding is stimulated) and of 150 V (structural unfolding takes place depending on the stability of the iron complex). Second, we measured the normalised collision energy necessary to fragment 50% of ferri-pyoverdine ions (CE50). This experiment was conducted on a Q Exactive hybrid quadrupole-Orbitrap mass spectrometer (Thermo Fisher Scientific, Waltham, MA, USA). Ferri-pyoverdine ions were isolated and fragmented at increasing normalised collision energies (NCE) using parallel reaction monitoring (PRM), while the ion intensity of the intact ion precursor was evaluated.

*Crude pyoverdine purification*

To stimulate pyoverdine production, 500 mL CAA medium was supplemented with 250 µM of the synthetic iron chelator 2,2’-bipyridyl to create an iron-limited environment and inoculated with 2 mL of washed overnight *Pseudomonas* cultures. We incubated the cultures at 28 °C for 120 hours with agitation (170 rpm). Cultures were then centrifuged at 15,049*g* for 15 minutes. We then harvested the supernatant and adjusted its pH to 6 using 1 M HCl. We ran the supernatants over Amberlite XAD-16N resin (Sigma Aldrich, Switzerland) columns at a rate of 2 drops per second and cleared the column of any salts by using 500 mL Milli-Q water. Pyoverdines were then eluted with 50% methanol and fractions of 50 mL were collected. Fractions containing the highest amount of pyoverdine (measured by fluorescence, excitation: 400 nm and emission: 460 nm), were pooled. After evaporation of the methanol, we lyophilized the samples for 24 hours and stored them at -20 °C. As the crude purified extracts may contain small impurities, the exact pyoverdine concentration is unknown and the concentrations. For the figures, concentrations were divided by 6 mg/mL (highest concentration used) and expressed as relative concentrations.

*HPLC purification of pyoverdines and ferribactin*

For HPLC purification, we first crudely purified the pyoverdine from 9 L supernatant as described above. The crude extract was then dissolved in water and fractionated by preparative HPLC at room temperature (Agilent 1260 Infinity system, equipped with a Phenomenex Luna 5 µm C18 21.2 x 250 mm column). Ultraviolet (UV) detection was carried out at λ = 210 nm, 230 nm, 280 nm, 350 nm, 400 nm, and 460 nm. Deionized water (Milli-Q, Millipore) (solvent A) and acetonitrile (solvent B) were used as the mobile phase with a flow rate of 15 mL/minute. We eluted with a gradient of 5% – 100% solvent B in 40 minutes, followed by isocratic conditions at 100% solvent B for 10 minutes. The fractions containing pyoverdine were identified based on the HR-ESI-MS data acquired on a Thermo Scientific Q Exactive hybrid quadrupole-orbitrap mass spectrometer (scan range 100–2500 *m/z*, capillary voltage 4500 V, dry temperature 200 °C) coupled to an UltiMate 3000 UHPLC system (Dionex, equipped with a Kinetex 2.6 µm XB-C18 150 x 4.6 mm column; solvent A: deionized water; solvent B: acetonitrile; gradient, 5% B for 30 seconds increasing to 100% B in 15.5 minutes and then maintaining 100% B for 5 minutes; flow rate 0.6 mL/minute; UV–Vis detection 200–600 nm). The fractions containing pyoverdine eluting at a retention time (RT) of 3.84 minutes in the LC-MS chromatogram (Figure S7) were further purified on a semi-preparative reverse phase HPLC at room temperature (Agilent 1260 Infinity system, equipped with a Phenomenex Luna 5 µm Phenyl-Hexyl 10 x 250 mm column, λ = 210 nm, 230 nm, 280 nm, 350 nm, 400 nm, and 460 nm monitored by UV detection). The elution gradient was increased from 5% to 15% solvent B for 30 minutes followed by a gradient shift from 15% - 100% in 5 minutes, and finally isocratic condition at 100% solvent B for 5 minutes. During the pyoverdine purification we noted high amounts of the precursor molecule ferribactin (RT of 2.42 minutes in the LC-MS chromatogram; Figure S8). We thus purified it alongside with pyoverdine using the same method.

*Cytotoxicity assays*

To determine the cytotoxicity of pyoverdine treatments against mammalian cells, we subjected human embryonic kidney 293 (HEK-293) and mouse motor neuron-like neuroblastoma-spinal cord hybrid (NSC-34) cells to the crude extract of the three most potent pyoverdines (3A06, 3G07, s3b09). HEK-293 cells (Invitrogen, R78007) were cultured in Dulbecco’s modified Eagle medium (DMEM; Sigma, D5671) supplemented with 10% fetal bovine serum (FBS; Gibco, 10270-106), 1X GlutaMAX (Gibco, 35050-061), 100 U/mL penicillin and 100 µg/mL streptomycin (Gibco, 15140-122). NSC-34 cells (Bioconcept, CLU140) were proliferated on Matrigel (Corning, 354234)-coated dishes in DMEM supplemented with 10% FBS, 1X GlutaMAX, 100 U/mL penicillin and 100 µg/mL streptomycin. For experiments, differentiation was induced by switching to DMEM/F12 medium (Gibco, 21331-020) supplemented with 1X GlutaMAX, 1X B27+ supplement (Gibco, 17504-044), 1X N2 supplement (Gibco, 17502-048), 20 ng/mL BDNF (PeproTech, 450-02), 20 ng/mL GDNF (PeptroTech, 450-10), 100 U/mL penicillin and 100 µg/mL streptomycin. All cells were cultured at 37 ºC with saturated humidity and an atmosphere of 5% CO_2_.

For the assay, 10,500 cells/well were plated on 96-well plates (Greiner Bio-One, 655090) and, after 48 hours, the culture medium was replaced and the pyoverdines were added at the appropriate concentrations. For this, pyoverdine stocks at 240 mg/mL were diluted in MilliQ water such that the addition of 5 µL of the pyoverdine solution to 195 µL of the culture medium resulted in final relative concentrations between 1 and 0.016. As the exact pyoverdine concentration in the crude extracts is unknown, the above concentrations were divided by the highest concentration used (6 mg/mL) and expressed as relative concentrations in the figure. After further 48 hours, the pyoverdine-containing culture medium was removed, and cells were incubated with 1 µg/mL thiazolyl blue tetrazolium bromide (Sigma, D5655) in 100 µL of fresh medium for 1.5 hours (NSC-34) or 40 minutes (HEK-293). The reaction was stopped, and formazan crystals simultaneously solubilized with the addition of 100 µl of a solution containing 10% sodium dodecyl sulfate (SDS; Sigma-Aldrich, 05030) and 0.03% HCl (Supelco, 100319). Finally, the absorbance of cell debris and other contaminants at 630 nm were subtracted from the absorbance of the solubilized formazan at 570 nm.

*Infection model*

All larvae were ordered from a local vendor (Bait Express GmbH, Basel, Switzerland) and stored without food at 16 °C in the dark for up to 3 days. In a first step, we determined the mean weight of 336 final instar *G. mellonella* larvae (mean = 435 mg). For all subsequent experiments, we weighed the larvae and only used individuals with a mean weight ± 20% (range: 340 mg - 530 mg). Larvae were immobilized on ice for 15 minutes, surface-sterilized with 70% ethanol and randomly distributed to individual wells on 24-well plates. 10 larvae were used per infection-treatment combination. To inject bacteria and to administer treatments, we used sterile needles (Sterican 26G, 0.45 x 12 mm (Braun)) and syringes (1 mL injekt-F (Braun)) attached to a syringe pump (New Era Pump Systems Inc, model NE-300) set to a flow rate of 5 mL/hour. For both infection and treatment delivery, a volume of 10 µL per larvae was injected between the posterior larval prolegs.

For infection experiments, we grew *A. baumannii*, *K. pneumoniae* and *P. aeruginosa* overnight as described above, washed them twice with PBS and diluted them to an OD_600_ of 1 in PBS. This OD_600_ corresponds to 4*10^8^ CFU/mL of *A. baumannii* and 9*10^8^ CFU/mL of *K. pneumoniae* and *P. aeruginosa*. We conducted a first experiment to determine the infection dose that kills approximately 50% of the larvae (LD_50_ value). For this, we diluted the cells in PBS and injected a range of bacterial dilutions into the larvae (in a volume of 10 µL, Figure S5). From now onwards, we express the infectious dose as CFU/larvae. For *A. baumannii*, we tested dilutions between 4*10^1^ - 4*10^6^ CFU/larvae and found that 1.8*10^5^ CFU/larvae corresponded to LD_50_. For *K. pneumoniae*, we tested dilutions between 9*10^1^ - 9*10^6^ CFU/larvae with 8.9*10^5^ CFU/larvae being closest to LD_50_. Finally, for *P. aeruginosa* we tested dilutions between 4 - 116 CFU/larvae and found that it was not possible to determine an LD_50_ value for this species, because all CFU/larvae doses lead to a 100% death rate (Figure S5).

*Experimental evolution experiment plate design and setup*

For each pathogen-treatment combination, we included six independently replicated lineages (populations) on 96-well plates. Each plate further contained two control populations that evolved in plain CAA medium, except for *K. pneumoniae*, for which all six controls were on the same plate. The six replicate populations were arranged along the diagonal of 96-well plates, or at the border of the plates, in such a way that they were surrounded by medium on all sides. This plate layout minimized the chance of cross-contamination and resulted in a total of ten plates (three each for *A. baumannii*, *P. aeruginosa* and *S. aureus* and one for *K. pneumoniae*).

Prior experimental evolution, we prepared batches of media for the entire duration of the experiment. Specifically, all treatments were prepared in CAA medium, then aliquoted into the respective wells of 96-well plates, and all plates were stored at -20 °C. At the start, we grew the pathogens overnight, washed and diluted them as described in the methods, and transferred 2 µL of the diluted pathogens into 200 µL medium in 96-well plates. Subsequently, the plates were incubated at 37 °C for 22 hours, shaken at 170 rpm. After incubation, we measured growth (OD_600_) of all evolving lineages with the plate reader. We then diluted the evolving cultures 1:1,000 in CAA medium and added them to a fresh treatment plate thawed from the freezer. After the transfer, the new plates were again incubated at 37 °C and 170 rpm agitation for 22 hours. We added glycerol to the old plates to a final concentration of 25% and stored them at -80 °C. We observed a steady decrease in growth in all *S. aureus* populations, suggesting that we overdiluted these cultures. Consequently, we reduced the dilution to 1:10 from day 4 onwards. Conversely, we observed a steady increase in growth for *A. baumannii* and *P. aeruginosa* populations, suggesting that these cultures were not diluted enough. Consequently, we increased dilution rates to appropriate levels on day 4, day 8, and day 12.

*Genomic analyses of evolved populations and clones*

For sequencing, we grew the clones and populations in 12 mL LB or TSB (*S. aureus*) at 37 °C and 170 rpm and harvested cultures upon reaching an OD_600_ between 0.8-1 by centrifugation (7,500 rcf, 3 minutes). We washed cultures in 0.8% NaCl and resuspended the pellet in DNA shield buffer (Zymo research). Samples were then sent to MicrobesNG (Birmingham, United Kingdom) for library preparation and sequencing. Whole-genome sequencing was performed on the Illumina NovaSeq6000 platform (Illumina, San Diego, USA) using a 250 bp paired end protocol. Adapter sequences were trimmed using Trimmomatic v0.30^3^ with quality cut-off of Q15. De novo assemblies were performed using SPAdes v3.7^4^ and contigs annotated with Prokka v1.11^5^.

*References*

1.	Butaitė, E., Baumgartner, M., Wyder, S. & Kümmerli, R. Siderophore cheating and cheating resistance shape competition for iron in soil and freshwater Pseudomonas communities. *Nat. Commun.* **8,** 414 (2017).

2.	Kramer, J., López Carrasco, M. Á. & Kümmerli, R. Positive linkage between bacterial social traits reveals that homogeneous rather than specialised behavioral repertoires prevail in natural Pseudomonas communities. *FEMS Microbiol. Ecol.* **96,** (2020).

3.	Bolger, A. M., Lohse, M. & Usadel, B. Trimmomatic: a flexible trimmer for Illumina sequence data. *Bioinformatics* **30,** 2114–2120 (2014).

4.	Bankevich, A., Nurk, S., Antipov, D., Gurevich, A., Dvorkin, M., Kulikov, A. S., Lesin, V. M., Nikolenko, S. I., Pham, S., Prjibelski, A. D., Pyshkin, A. V., Sirotkin, A., Vyahhi, N., Tesler, G., Alekseyev, M. A. & Pevzner, P. A. SPAdes: a new genome assembly algorithm and its applications to single-cell sequencing. *J. Comput. Biol.* (2012).

5.	Seemann, T. Prokka: rapid prokaryotic genome annotation. *Bioinformatics* **30,** 2068–2069 (2014).

**Supporting Figures**

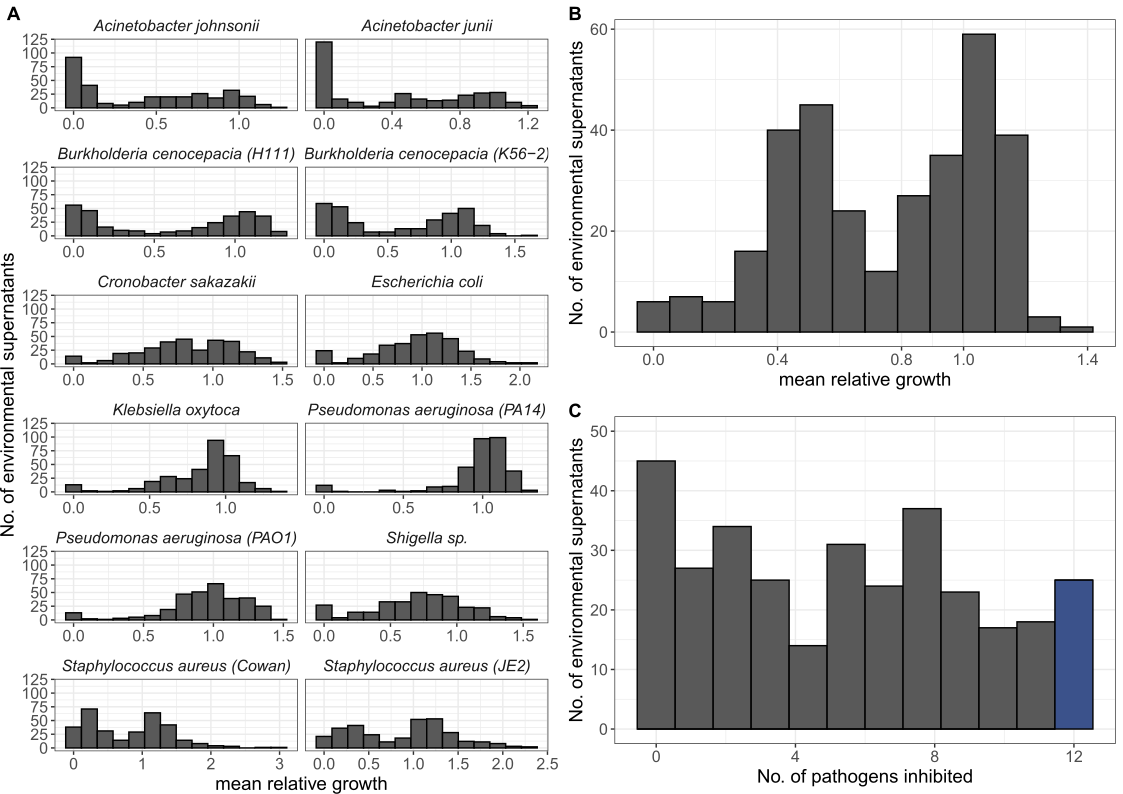
**Figure S1 | Growth of pathogens in supernatant treatments and the number of pathogens inhibited by a supernatant**. (A) Histograms showing the distribution of relative growth (mean across n=4 replicates) for each of 12 human opportunistic pathogens when exposed to the supernatants of the 320 natural *Pseudomonas* spp. (B) Histogram showing the distribution of the mean relative growth across all 12 pathogens when exposed to the 320 supernatants. (C) Histogram showing the number of pathogens that were reduced in their growth by a specific supernatant (at least 20% growth reduction).

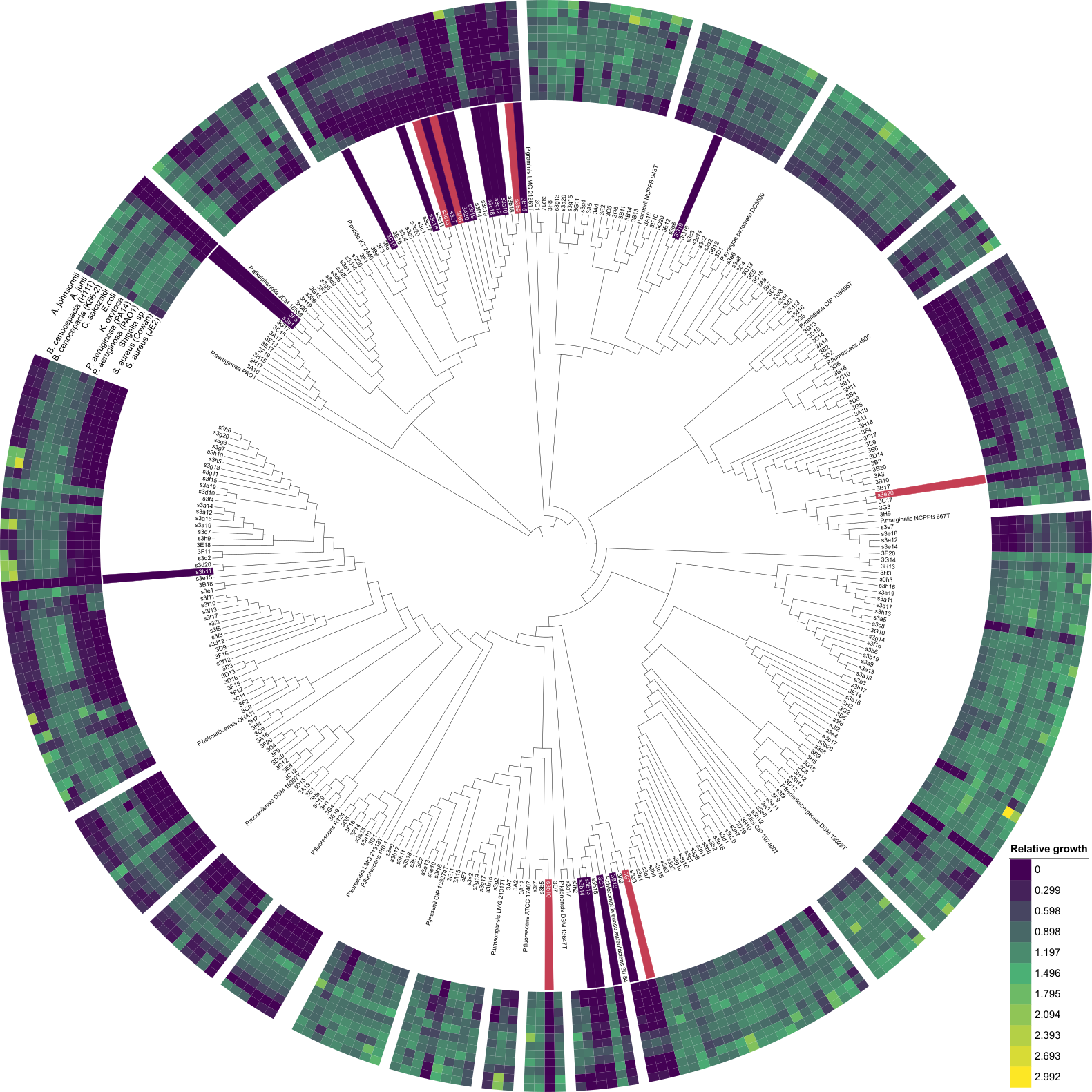

**Figure S2 | Cladogram of environmental *Pseudomonas* isolates from soil and pond habitats based on partial *rpoD* sequences.** The tree includes 297 out of the 320 *Pseudomonas* isolates for which *rpoD* sequences with lengths ≥600 bp were available. 23 isolates with shorter sequence lengths had to be excluded. *rpoD* sequences stem from Butaitė et al. 2018 <https://doi.org/10.1111/1462-2920.14355> and are available through the European Nucleotide Archive (ENA, accession number: PRJEB21289). *P. aeruginosa* PAO1 was used as the outgroup. *rpoD* sequences from 20 well-characterised fluorescent pseudomonads were taken from the literature and integrated into the cladogram to cover taxonomic affiliations of our strains. Concentrical rings depict the supernatant-mediated growth effects of the environmental isolates on the 12 pathogens as a heatmap (matching the results from Fig. 1), ranging from complete growth inhibition (dark purple) to growth promotion (yellow). Isolates whose supernatants strongly inhibited all 12 pathogens are marked in dark purple, while the top candidates, which inhibited those pathogens exclusively under iron-limited conditions, are marked in red (*rpoD* sequences available for 23 out of these 25 strains).

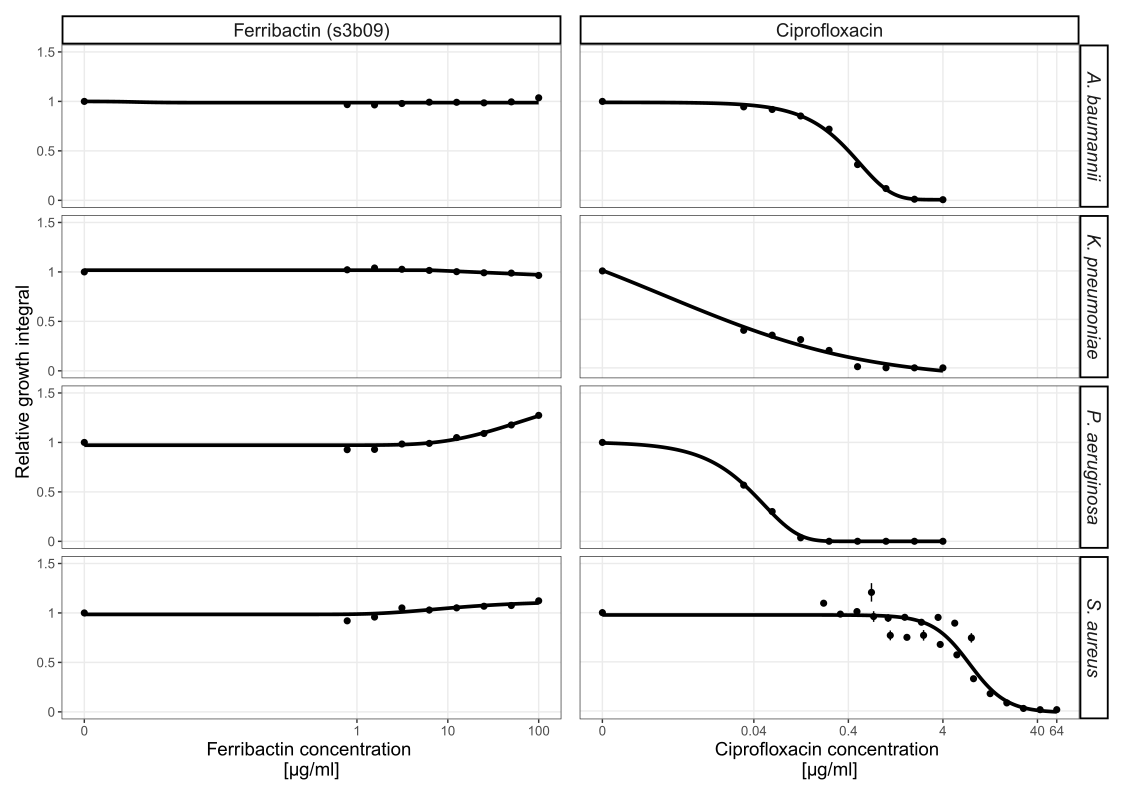

**Figure S3 | Ferribactin and ciprofloxacin dose-response curves for *A. baumannii*, *K. pneumoniae*, *P. aeruginosa* and *S. aureus.*** We exposed the four human opportunistic pathogens to a concentration gradient of HPLC-purified ferribactin s3b09 and ciprofloxacin. Growth values are scaled relative to the untreated control in untreated CAA medium. Dots and error bars show mean values and standard errors, respectively, across 3 replicates. Dose-response curves were fitted using 4- or 5-parameter logistic regressions.

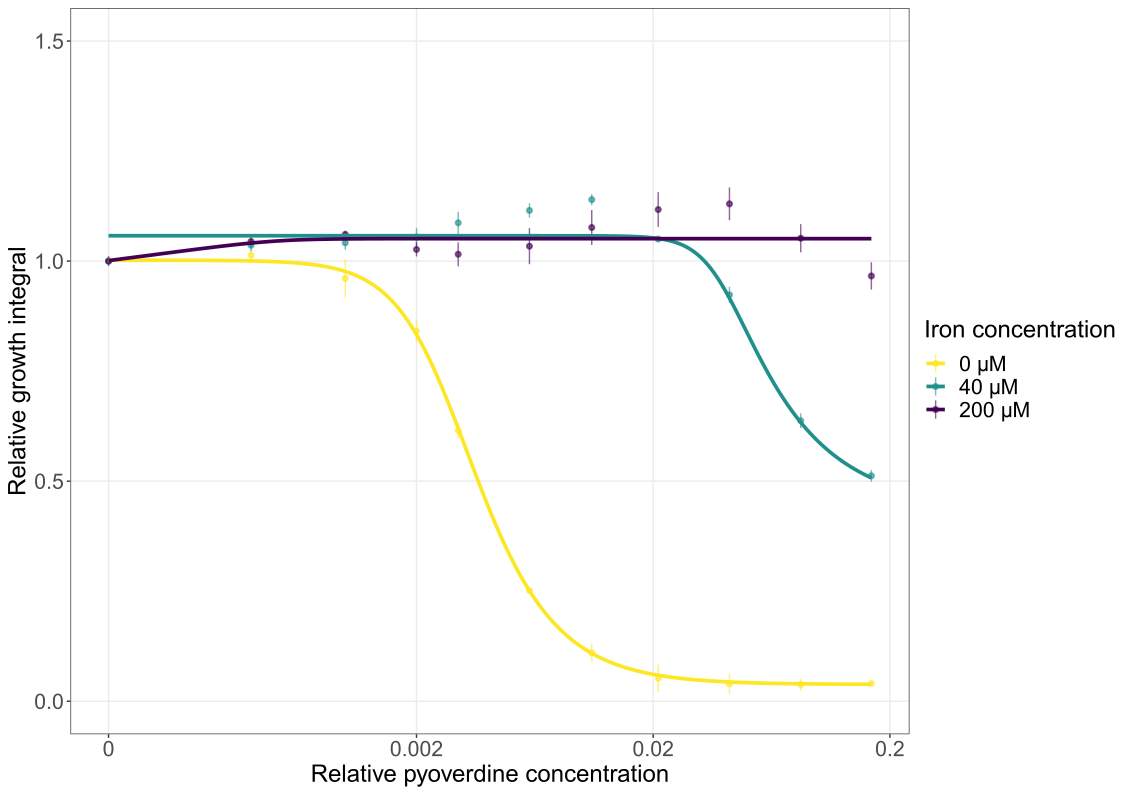

**Figure S4 | The effect of iron-saturated pyoverdines on the growth of *A. baumannii*.** We exposed *A. baumannii* to increasing crude pyoverdine s3b09 concentrations (relative concentrations 0.16 to 0.0004) in CAA medium (yellow line, n = 5), or in CAA medium supplemented with either 40 µM (green line, n = 3) or 200 µM FeCl_3_ (purple line, n = 3). Growth values are scaled relative to the pyoverdine untreated control, independently for each of the three media (CAA with either 0, 40 or 200 µM FeCl_3_). Dots and error bars show mean values and standard errors, respectively. Dose-response curves were fitted using 4- or 5-parameter logistic regressions.

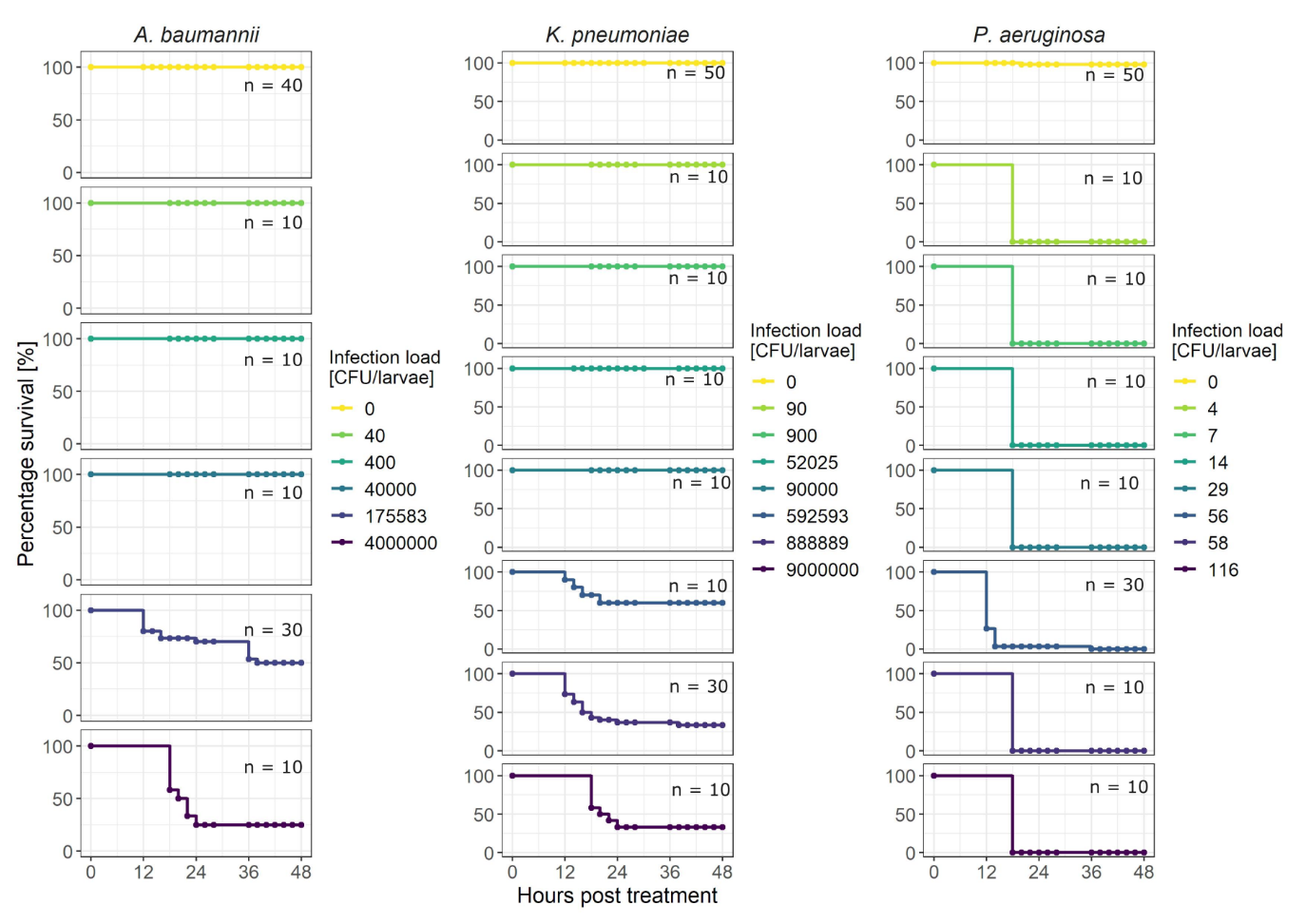

**Figure S5 | Survival curves of *G. mellonella* infected with *A. baumannii*, *K. pneumoniae* or *P. aeruginosa* at different infection loads.** To determine the infection load at which approximately 50% of larvae die (LD_50_) after 48 hours, we infected larvae with different infection loads with one of three human opportunistic pathogens, *A. baumannii*, *K. pneumoniae*, or *P. aeruginosa*. **All panels show the** percentage of survival of wax moth larvae over 48 hours post infection. Based on the resulting killing curves, we selected 1.8 * 10^5^ CFU/larvae for *A. baumannii* and 8.9*10^5^ CFU/larvae for *K. pneumoniae* for the main infection and treatment experiments. These values correspond closely to LD_50_. In contrast, no LD_50_ value could be determined for *P. aeruginosa* as 100% of the larvae died, regardless of the infection dose. For the main experiments, we took 56 CFU/larvae.

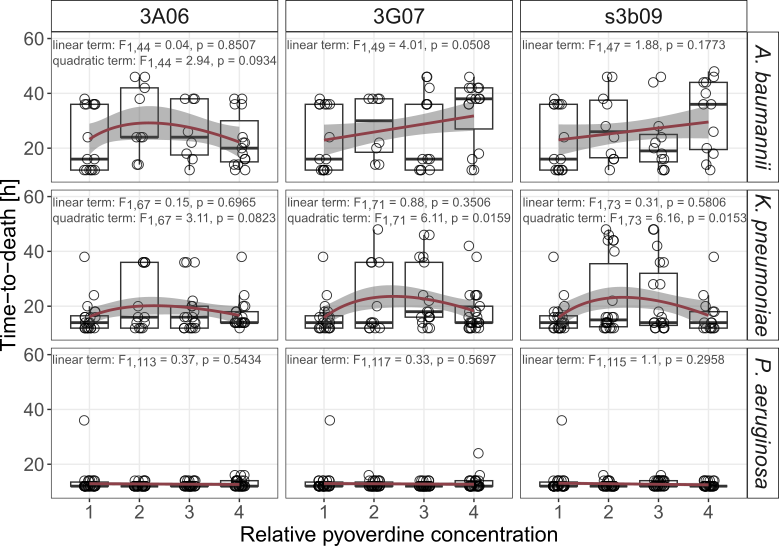

**Figure S6 | Variation in the time-to-death of *G. mellonella* larvae in response to pyoverdine treatment.** Larvae were infected with either *A. baumannii*, *K. pneumoniae* or *P. aeruginosa* and treated with either pyoverdine 3A06, 3G07 or s3b09 at relative pyoverdine concentrations 0, 0.01, 0.05 or 0.1. Data stem from three independent experiments, each with 10 larvae per infection and treatment. For this analysis, only larvae that died during the experiment could be included. Box plots show the median and the first and third quartiles, and whiskers represent the 1.5x interquartile range. Red lines and shaded areas show the linear or quadratic regression lines (based on square-root transformed data) and 95% confidence intervals, respectively.

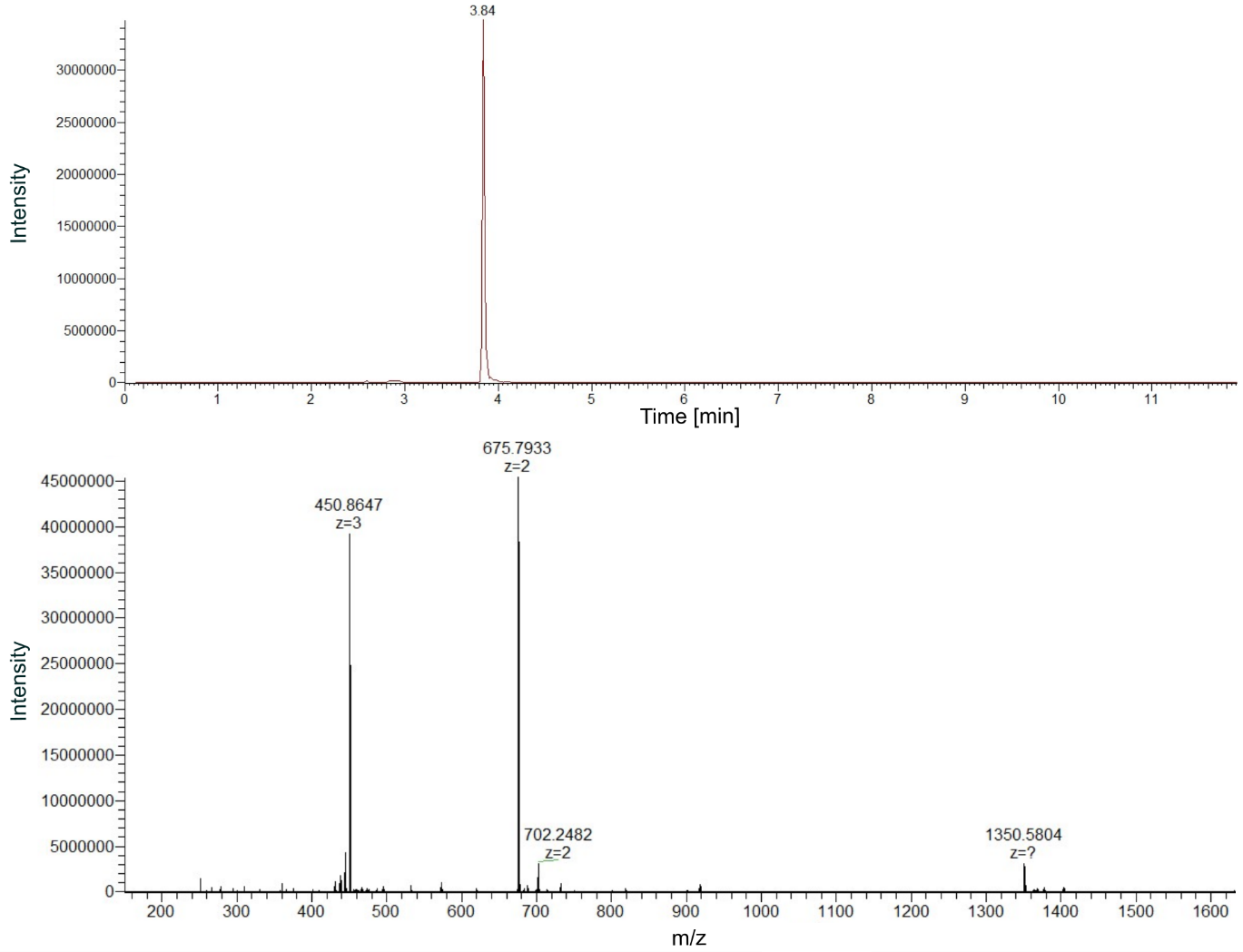

**Figure S7 |** **High resolution mass spectrum (HRMS) of purified pyoverdine s3b09.** Mass chromatogram (upper panel) and spectrum (lower panel) of the HPLC-purified pyoverdine s3b09. The compound elutes at a retention time of 3.84 minutes. It is the compound with the highest abundance in the measured sample. Based on these data, we define the measured sample as highly purified pyoverdine.

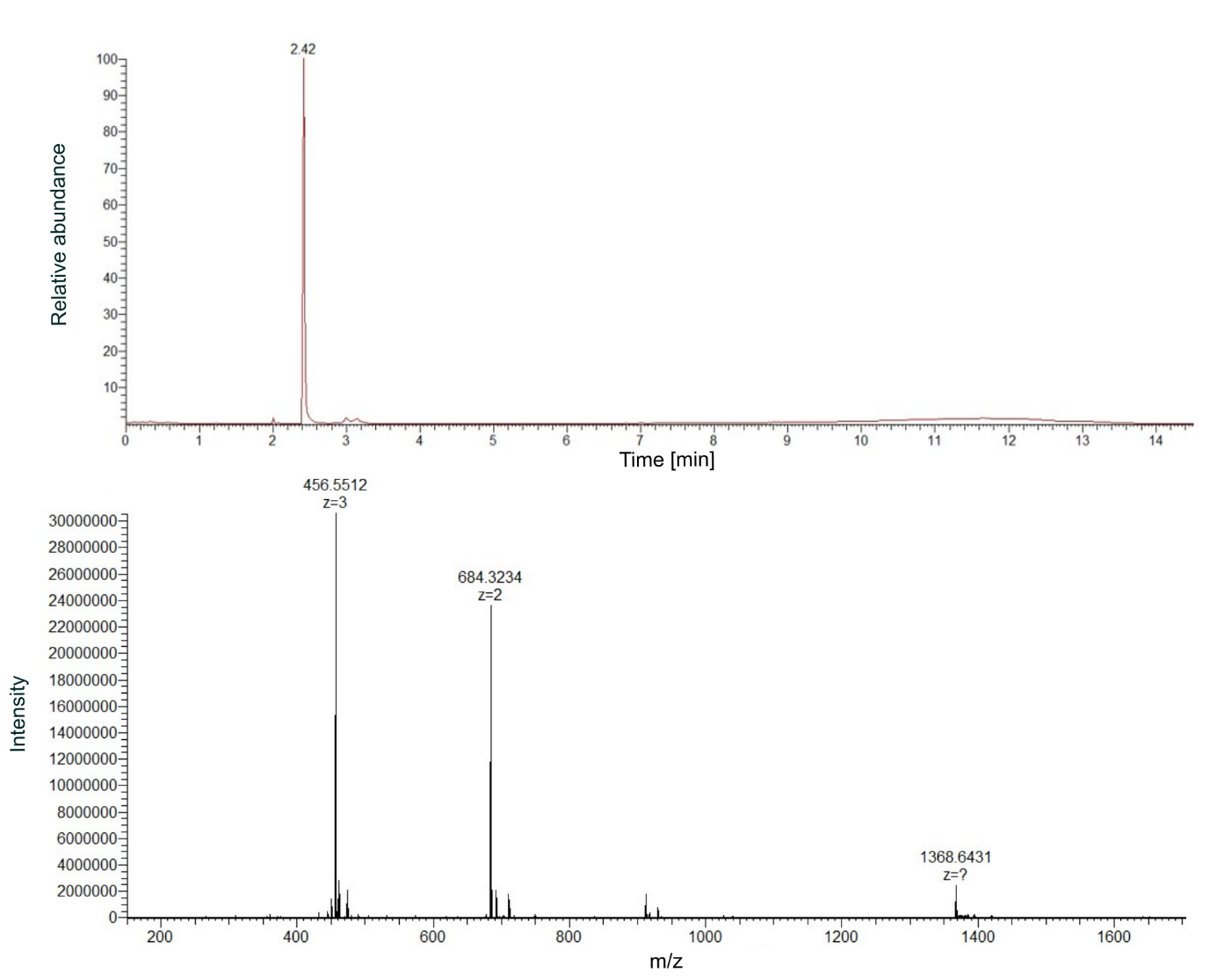

**Figure S8 |** **High resolution mass spectrum (HRMS) of purified ferribactin (precursor molecule of pyoverdine s3b09).** Mass chromatogram (upper panel) and spectrum (lower panel) of ferribactin. The compound elutes at a retention time of 2.42 minutes and has the highest intensity in the measured sample.

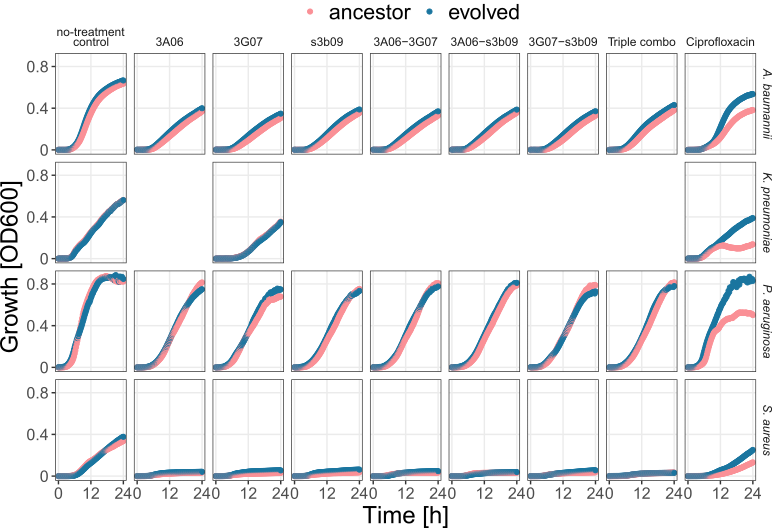

**Figure S9 | Population growth kinetics of the pathogens *A. baumannii*, *K. pneumoniae*, *P. aeruginosa* and *S. aures* before and after experimental evolution.** Growth kinetics (measured at OD 600nm) of ancestral (pink) and evolved (blue) populations of the four pathogens subjected to a no-treatment control, single pyoverdine 3A06, 3G07 or s3b09 treatments, their double and triple combination treatments, and a ciprofloxacin antibiotic treatment.

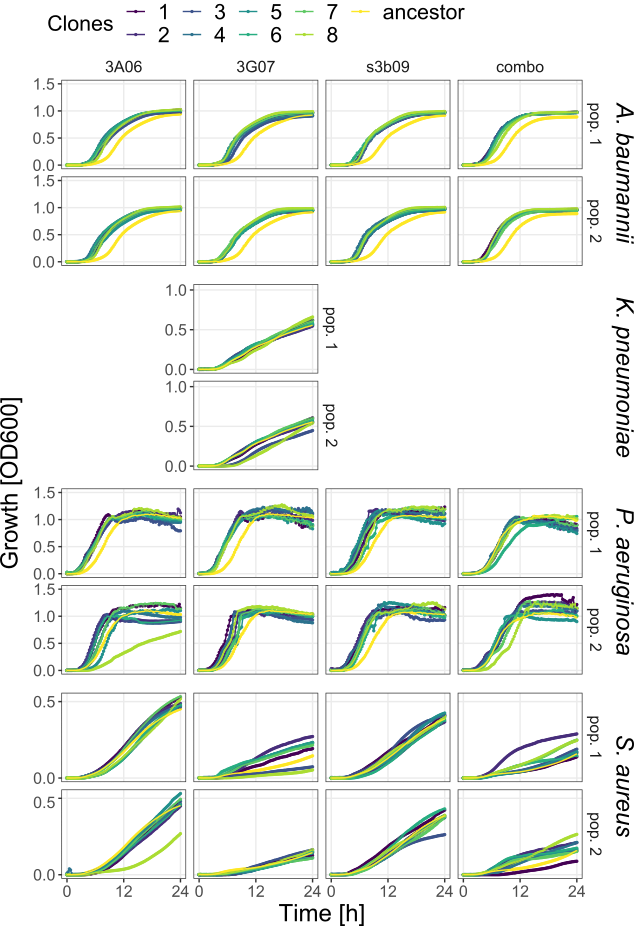

**Figure S10 | Growth kinetics of evolved clones of the pathogens *A. baumannii*, *K. pneumoniae*, *P. aeruginosa* and *S. aureus* in comparison to the ancestor.** We exposed the ancestor (yellow) and eight evolved clones per treatment (pyoverdines: 3A06, 3G07, s3b09, combo) for two lineages (pop.1 and pop.2; 208 individual clones in total) to the treatment they evolved in and measured their growth (OD at 600nm) over time.

**Table S1 | Survival analysis for *G. mellonella* larvae infected with bacterial pathogens and treated with pyoverdine.** We calculated the hazard ratios and Wald statistics for Cox proportional hazard regressions for the survival of *G. mellonella* larvae following infection with *A. baumannii*, *K. pneumoniae* or *P. aeruginosa*: For pyoverdine-only treatments, we used the log-rank test and adjusted for multiple comparisons using the Holm method. P-values below 0.05 were considered significant.

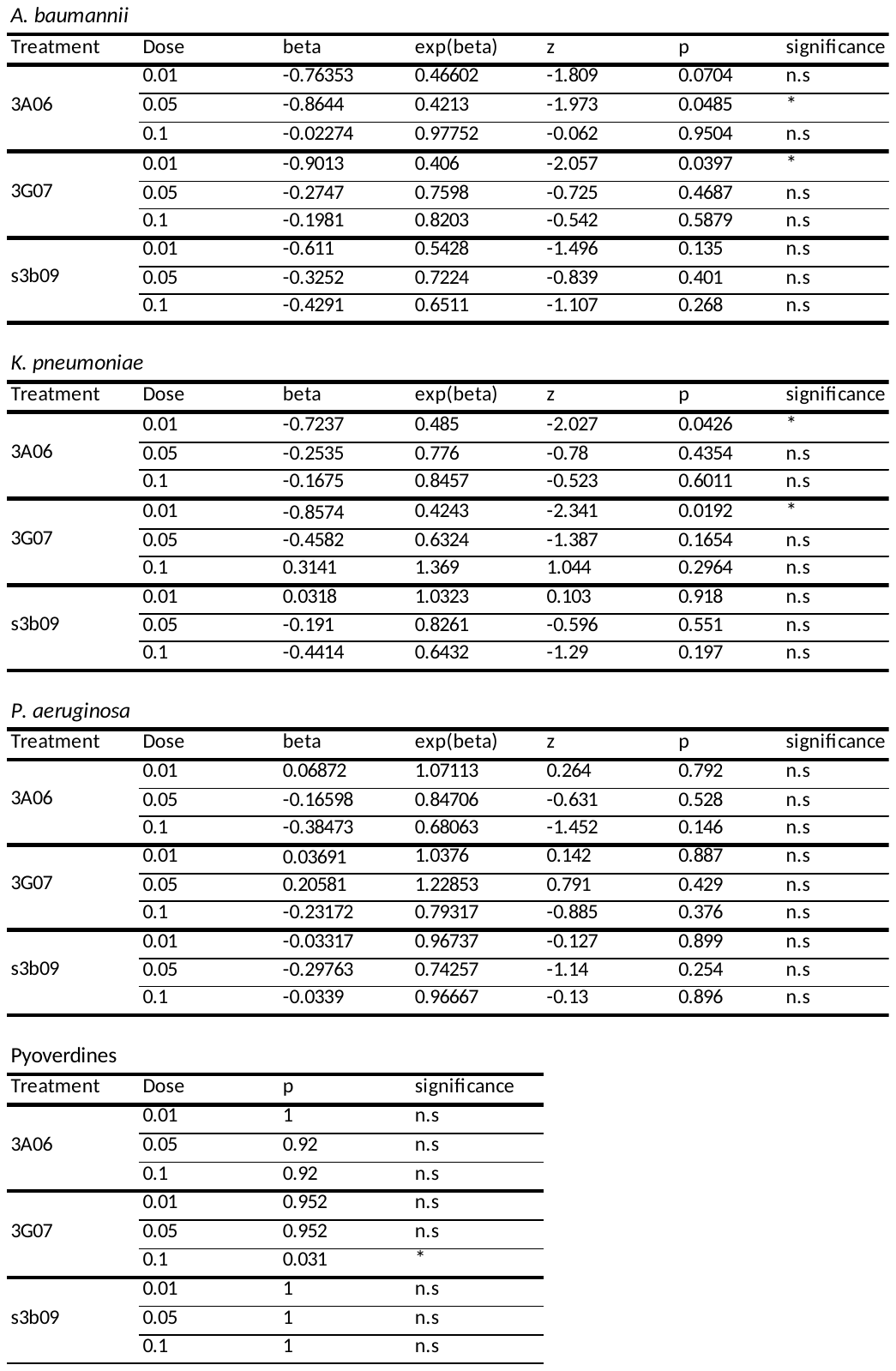

**Table S2 | Single (crude) pyoverdine and ciprofloxacin concentrations used for experimental evolution**. Values represent rounded IC50 concentrations inferred from the dose-response curves shown in Figure 3. The pyoverdine concentrations are expressed as relative concentrations (to the highest concentration used), while ciprofloxacin concentration is given in µg/mL. For *P. aeruginosa*, no IC50 concentrations could be calculated for pyoverdines 3A06 and s3b09 and the values therefore represent the concentrations for which the bacterial growth was lowest.

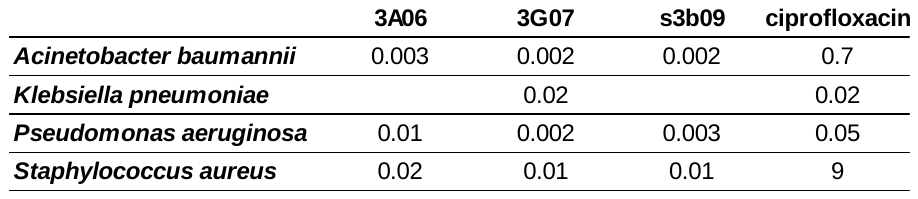

**Table S3 | Statistical analyses from the population and clone screening after experimental evolution**

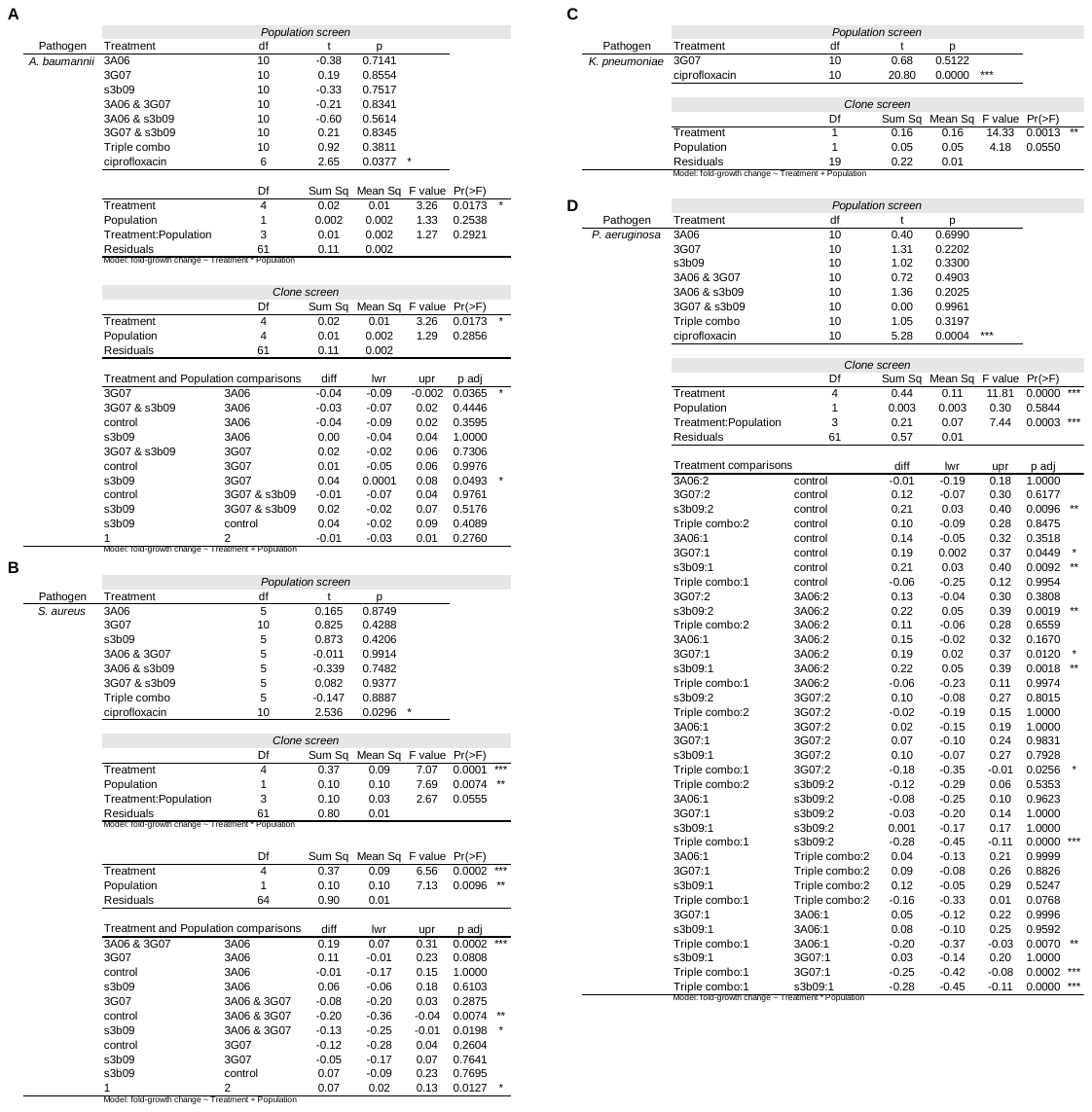

**Table S4 | List of mutations per species and clone that were not associated with adaptation to medium.**

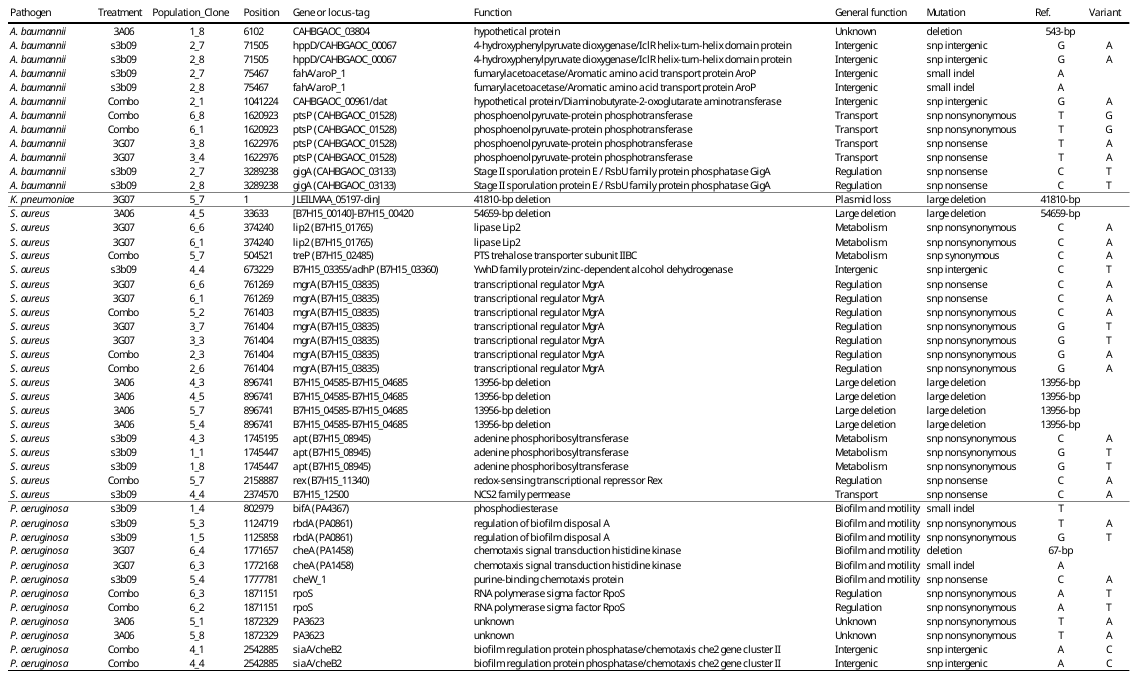

**Table S5 | Large deletions (≥ 600 bp) in *K. pneumonia*e and *S. aureus* clones.**

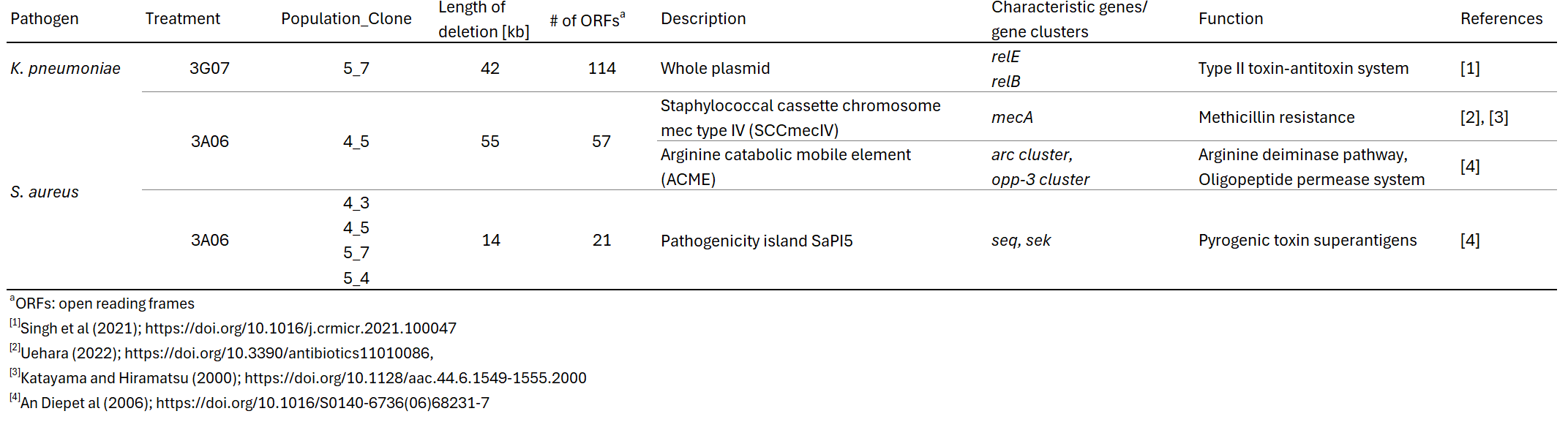
